# LSD1 serine 166 is a phosphorylation switch for chromatin landscaping, gene activation, and tissue remodeling

**DOI:** 10.1101/2025.05.14.653937

**Authors:** Mukund Sharma, Anna Aakula, Oxana V. Denisova, Elena Ceccacci, Kourosh Hayatigolkhatmi, Jonatan Caroli, Reetta Nätkin, Marie-Catherine Laisne, Riikka Huhtaniemi, Sujan Ghimire, Johannes Merilahti, Roberta Noberini, Jesse Kamila, Otto Kauko, Andrea Mattevi, Tiziana Bonaldi, Guillaume Jacquemet, Matti Nykter, Saverio Minucci, Jukka Westermarck

**Affiliations:** Turku Bioscience Centre, University of Turku and Åbo Akademi University, 20520 Turku, Finland; InFLAMES Research Flagship Center, University of Turku and Åbo Akademi University, 20520 Turku, Finland; European Institute of Oncology (IEO) IRCCS, Via Adamello 16, 20139 Milan, Italy; Department of Biology and Biotechnology, Via Ferrata 9, University of Pavia, Italy; Faculty of Science and Engineering, Cell Biology, Åbo Akademi University, 20520 Turku, Finland; Laboratory of Computational Biology, Faculty of Medicine and Health Technology, Tampere University and Tays Cancer Centre, Tampere, Finland; Department of Oncology and Hematology-Oncology, University of Milan, Milan, 20122 Italy; Institute of Biomedicine, University of Turku, Turku, Finland

**Keywords:** KDM1A, H3K4, H3K9, B56, RRBS, ATAC sequencing, mass spectrometry proteomics, extracellular matrix

## Abstract

LSD1 is a histone 3 (H3) demethylase that can either repress or activate gene expression. We discover here that the so far enigmatic balance between these two activities in non-hormonal cancer cells is regulated by phosphorylation of serine 166 (S166) on LSD1. SET-mediated Protein Phosphatase 2A (PP2A) inhibition in KRAS mutant cells promotes S166 phosphorylation. Endogenous LSD1 S166 alanine mutant (S166A) cells display H3 lysine 9 demethylation and acetylation, euchromatin, and gene activation. Mechanistically this is explained by the impaired interaction of S166A mutant LSD1 with repressor proteins SNAI2 and MYBP1. Functionally LSD1 S166A mutant cells display augmented beta1 integrin activity and stress fiber formation, and the mutant xenograft tumors have altered tumor microenvironment associated with increased macrophage recruitment. Collectively, PP2A-regulated S166 of LSD1 is a phosphorylation switch for epigenetic gene activation in non-hormonal cancer cells. Conceptually we demonstrate how dephosphorylation of one amino acid on a non-histone protein shapes chromatin landscape in cancer cells, and modify tumor stroma, and immune cell content.

**Highlights:**

• Mechanism for gene activation by LSD1 in non-hormonal cancers
• Single phosphorylation switch in a non-histone protein controls epigenetic landscape
• Epigenetic protein phosphorylation in cancer cells shapes tumour immune microenvironment
• Novel function for Protein Phosphatase 2A (PP2A) in epigenome regulation via LSD1

**Graphical abstract:** 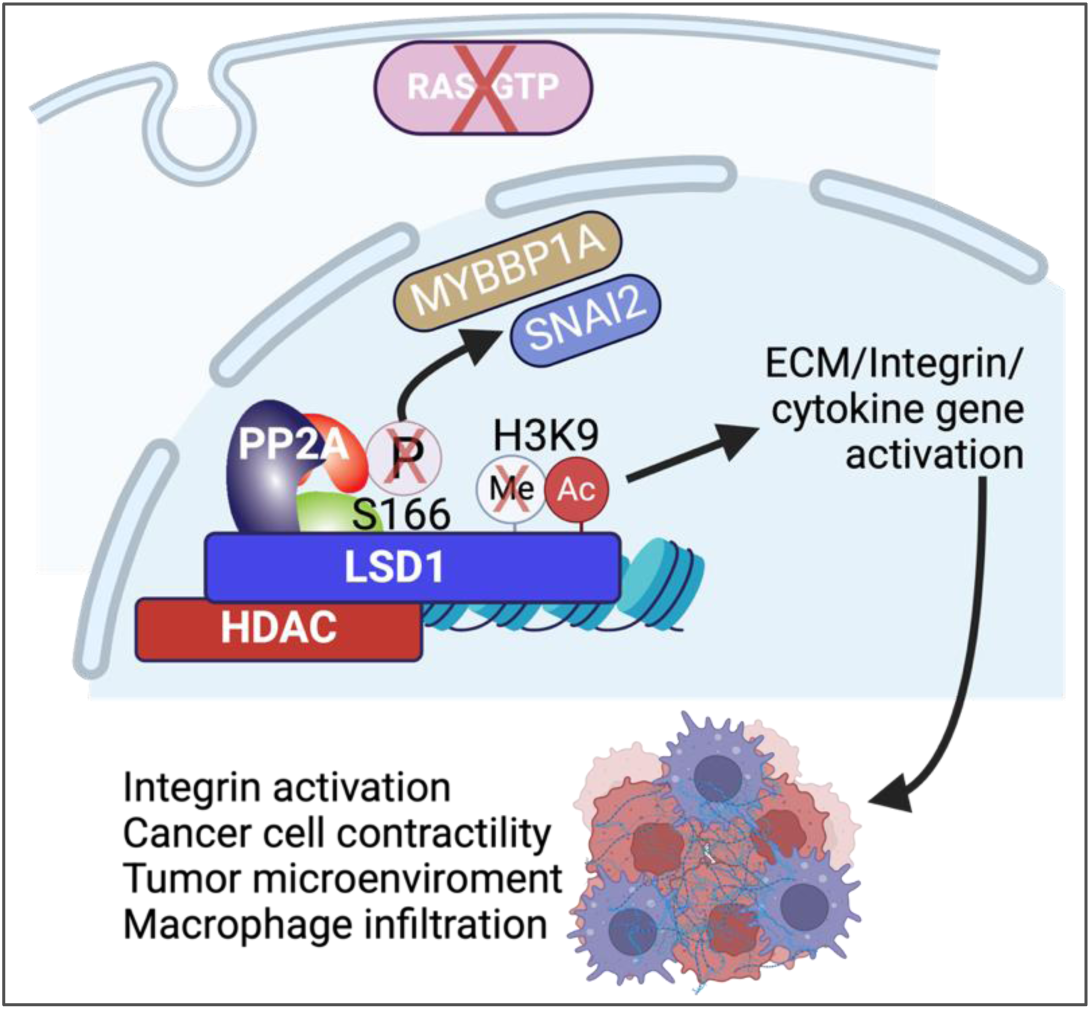

## Introduction

Gene expression is epigenetically controlled by chromatin accessibility of gene regulatory protein complexes and DNA methylation. These processes are regulated by number of epigenetic protein complexes facilitating either gene activation or repression (Laugesen and Helin, 2014). Reversible post-translational modifications (PTMs) of proteins allows cells in tissues to adapt to changing environments in seconds to minutes, whereas epigenetic mechanisms are considered to constitute part of cellular memory due to long-term impact on chromatin architecture. How these temporally distinct signalling systems are connected and convey changes from cellular environment to epigenetic gene regulation is however poorly understood (Separovich et al., 2020).

Among the PTMs, serine (S) phosphorylation is the most prevalent protein modification (Hornbeck et al., 2019) regulated by the balance between phosphorylating serine/threonine (S/T) kinases and dephosphorylating S/T phosphatases (Cordeiro et al., 2017; Hunter, 2012). Recent developments in mass spectrometry (MS) proteomics technologies have allowed comprehensive cataloguing of protein phosphorylation in different cell types and conditions; in fact, the current count of identified phosphorylation sites approaches 300 000 individual observations (Hornbeck et al., 2019). However, only a very small fraction (appr. 1-3 %) of identified phosphorylation events have so far been linked to any defined cellular function. Thereby, there are fundamental gaps in our basic understanding of how individual phosphorylation sites contribute to health and diseases (Hunter, 2012).

Among the most important mechanisms regulating cellular S/T phosphorylation, trimeric Protein Phosphatase 2A (PP2A) complexes are estimated to be able to dephosphorylate up to 70% of phosphorylated S/T amino acids (Fowle et al., 2019). In cancer cells, PP2A activity is regulated by overexpression of proteins such as SET, PME-1 or CIP2A (Kauko and Westermarck, 2018) that selectively inhibit B56 regulatory subunit containing PP2A trimers. Recently PP2A complexes, including SET-inhibited PP2A-B56, have been strongly implicated in transcriptional gene activation via their direct effects on RNA polymerase II C-terminal tail phosphorylation (Vervoort et al., 2021; Xu et al., 2024), but epigenetic protein targets for PP2A-B56 are poorly characterized. One of the important signaling mechanism regulated by PP2A-B56 is RAS signaling. RAS proteins are upstream activators of many serine/threonine kinase pathways (Klomp et al., 2024b), resulting ultimately in robust changes in the transcriptional outcomes (Klomp et al., 2024a). PP2A-B56 counteracts RAS activity in human cell transformation (Hahn et al., 2002; Sablina et al., 2010), and in various serine/threonine dependent signaling pathways (Westermarck and Hahn, 2008), but the extent of their shared target phosphoproteome has been unclear.

Lysine-specific histone demethylase 1A (LSD1)(a.k.a KDM1A) and its homologue LSD2 were the first enzymes shown to catalyse demethylation of histones (Ciccone et al., 2009; Perillo et al., 2020). Specifically, LSD1 demethylates either mono -or di-methylated histone 3 lysine 4 (H3K4me1/2) by its monoamine oxidase activity, that depends on the number of residues in the substrate-binding site and at the interface of the AOL-SWIRM domain (Perillo et al., 2020). Interaction of LSD1 with REST family of corepressor protein CoREST1 is required for LSD1-mediated for H3K4 demethylation (Yang et al., 2006). Importantly, in addition to its direct catalytic activity towards H3K4, LSD1 is also a critical member of the NuRD repression complex including histone deacetylases (HDACs). LSD1 also interacts with SNAIL family of zinc finger proteins that repress transcription. SNAI2 interaction also affects LSD1 target gene selectivity as it is known to impact especially genes that are involved in epithelial mesenchymal transition and cadherin regulation (Egolf et al., 2019). Thereby, protein interactions are in key role in explaining the functional diversity of LSD1 in gene repression (Perillo et al., 2020). Few kinases have been shown to phosphorylate LSD1 sites influencing its protein stability, protein interactions, and regulation of specific gene promoters, but regulation of LSD1 phosphorylation by phosphatases is very poorly understood (Karakatsanis et al., 2024). In hormonal cancers LSD1 has also gene activator function due to protein interactions with androgen receptor (AR) and estrogen receptor (ER)(Metzger et al., 2005; Perillo et al., 2020). However, whether LSD1 can function in gene activation in non-hormonal cancers has been debated, and this is a critical unanswered question with broad relevance.

Recently we profiled RAS and PP2A-regulated phosphoproteomes by siRNA-mediated inhibition of all major RAS isoforms, and by siRNA-mediated depletion of the PP2A-A subunit, or SET, PME-1, and CIP2A (Kauko et al., 2020; Kauko et al., 2015). Notably, analysis of the PP2A and RAS co-regulated phosphosites revealed significant enrichment in targets related to epigenetic gene regulation (Aakula et al., 2023). Chromatin organization and chromatin modifying proteins were strongly enriched as RAS-regulated processes also in a recent comprehensive phosphoproteome study (Klomp et al., 2024b). However, functional outcomes of PP2A -and RAS-mediated regulation of epigenetic complexes are poorly characterized. Here we demonstrate how single phosphorylation convergence site for PP2A and RAS signaling, serine 166 of LSD1, functions as a molecular switch for epigenetic landscaping, gene activation and tissue homeostasis. The results reveal both a mechanism for gene activation by LSD1 in non-hormonal cancers, but also generally indicate for an enormous regulatory potential of phosphorylation regulation of epigenetic proteins as the interface between phosphoproteome regulation and epigenetic gene regulation. On a conceptual level, the results indicate that systematic charting of phoshorylation sites at unstructured protein regions could fundamentally change our comprehension of non-genetic control in cancer and other pathological and physiological conditions.

## Results

### Identification of serine 166 of LSD1 as a direct target of PP2A/B56

We recently mapped the global impact of PP2A and RAS on cellular phosphorylation by mass spectrometry (MS)-based phosphoproteomics analysis in HeLa cells (Kauko et al., 2020; Kauko et al., 2015). This led to identification of dozens of previously uncharacterized putative PP2A and RAS target phosphorylation sites in epigenetic proteins (Aakula et al., 2023). One of these phosphorylation sites was serine 166 (S166) of LSD1 (Fig. 1A), that was found significantly increased in HeLa cells depleted of the PP2A scaffolding A subunit (PPP2R1A), resulting in global PP2A inhibition, whereas the site was dephosphorylated either by inhibition of PP2A inhibitor protein SET (i.e. by PP2A reactivation), or by combined depletion of H/N/K-RAS (Fig. 1A). Phosphorylation of S166 of LSD1 has been observed in more than 40 high content phosphoproteomics studies (Phosphositeplus), and across all 11 cancer types analysed in a recent pan-cancer study (Geffen et al., 2023). Among the LSD1 phosphosites observed in high throughput proteomics studies, S166 shows highest degree of evolutionary conservation (Fig. S1A). Further, phosphorylation of S166 of LSD1 was found vary across experimental conditions and cell lines in response to different stimuli based on the qPTM database (Yu et al., 2023)(Fig. 1B). However, no function has been assigned to this phosphosite yet.

**Figure 1.**
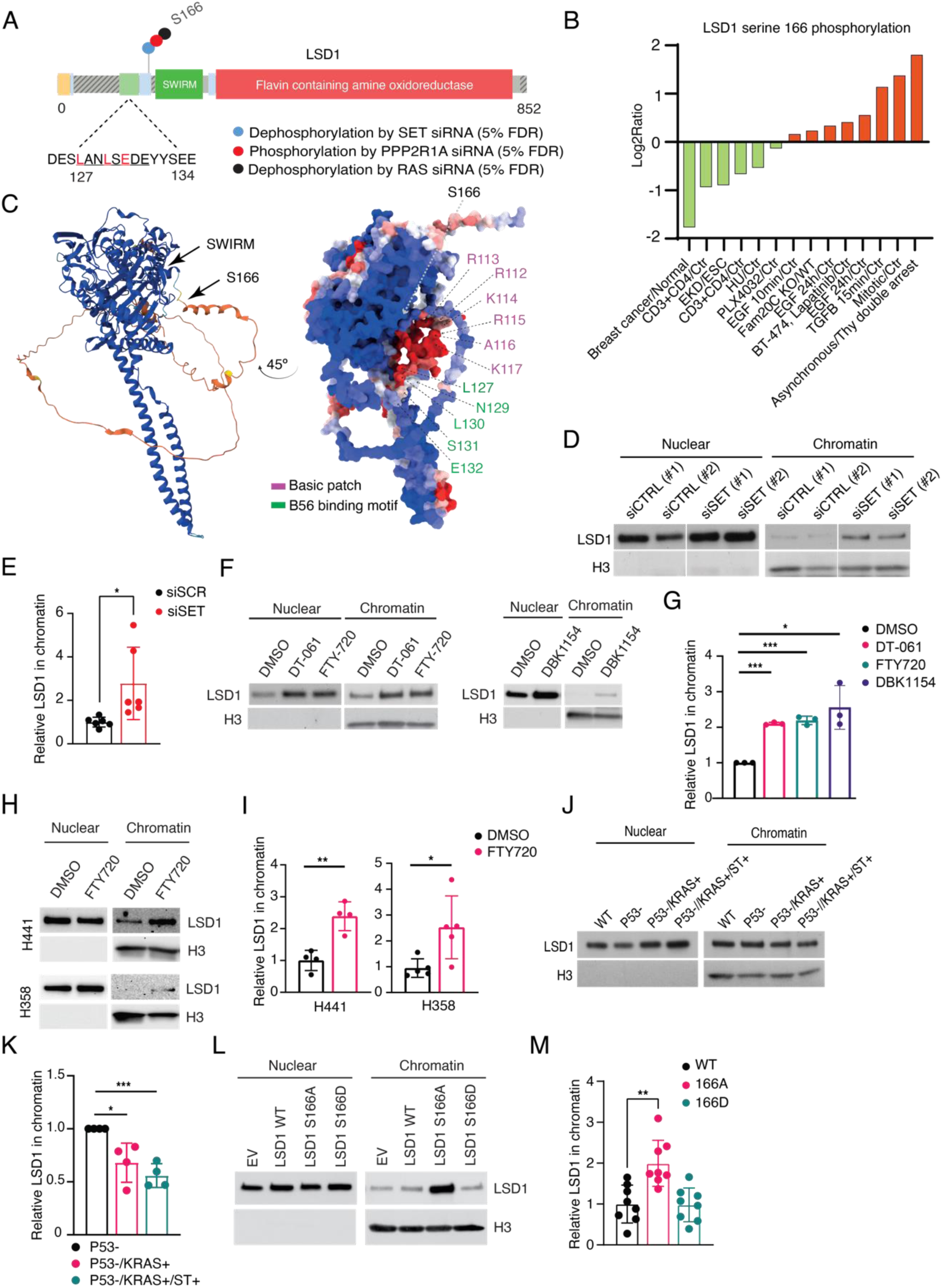
LSD1 serine 166 is a direct target for PP2A-B56. **(A)** Schematic presentation of RAS/PP2A-regulated phosphosite serine 166 (S166) on LSD1 indicated by a colored dot representing the change in phosphorylation by mass spectrometry upon siRNA treatments of either the PP2A inhibitory protein SET, PP2A-A scaffold protein PPP2R1A, or combined depletion of RAS isoforms (H/K/N). The amino acid sequence adjacent to the phosphosite is the consensus PP2A-B56 recognition motif. **(B)** Bar graph showing the phosphorylation dynamics of LSD1 at S166 across different experimental conditions and cell lines based on qPTM database. The log2 ratio represents phosphorylation enrichment or depletion. **(C)** Left panel: Overall structure of LSD1 from alphafold (AF-O60341-F1-v4) indicating location of S166 on the unstructured region. Right panel: LSD1 structure from alphafold colored by ScanNet binding site probability (from blue=low to red=high). LSD1 is rotated by 45° around the X-axis to provide an optimal view of the amino acids contributing to the basic patch (R113-K117) and B56 binding motif (L127-E132). **(D)** Western blot analysis of chromatin recruitment of LSD1 upon siRNA-mediated inhibition of the PP2A inhibitory protein SET in H460 cells. **(E)** Quantification of results from (D) (n = 3). **(F)** Western blot showing the impact of pharmacological PP2A activation (DT061, DBK1154, and FTY720) on LSD1 recruitment to chromatin in H460 cells. Each drug was used at 10 μM concentration for 48 h. **(G)** Quantification of results from (F) (n = 3). **(H)** Western blot showing the impact of pharmacological PP2A activation (FTY720) on LSD1 recruitment to chromatin in H358 and H441 cells. Each drug was used at 10 μM concentration for 24 h. **(I)** Quantification of results from (H) (n = 3). **(J)** Chromatin recruitment of LSD1 in the step-wise–transformed human bronchial epithelial cells. **(K)** Quantification of (J) (n = 4). **(L)** Western blot analysis of chromatin recruitment of LSD1 in H460 cells, transiently transfected with either wild type or S166A/D mutants of LSD1. **(M)** Quantification of results from (L) (n = 8). **(E, G, I, K, M)** Statistical Analysis: Data were analyzed using Student’s t-test, and results are presented as mean ± standard deviation (SD). Statistical significance was determined using the following thresholds: *p < 0.05, **p < 0.01, and ***p < 0.001.

Indicating that LSD1 is a direct PP2A/B56 complex substrate, we identified N-terminally from S166 a classical recognition sequence for the PP2A B56 subunit (127LANLSEDE134)(Hertz et al., 2016) (Fig. 1A). Upon structural and sequence-based disorder propensity analysis (Fig. 1A,C and S1B), S166 is one of the first amino acids of the N-terminal unstructured region recently demonstrated to be important for LSD1 function (Waterbury et al., 2024), but also adjacent to the functionally important SWIRM domain that is important for histone binding (Fig. 1A,C). Surface electrostatic potential analysis of LSD1 further revealed a cluster of basic residues adjacent to LSPI motif, and B56 affinity to its target proteins is known to be increased by such basic patch (Fig. 1C)(Wang et al., 2020). We confirmed interaction between LSD1 and the B56a regulatory subunit of PP2A by co-immunoprecipitation analyses (Fig. S1C, D). Serine 131 of the LSD1 LSPI motif corresponds to a phosphorylation site shown to increase B56 affinity to other B56 targets (Hertz et al., 2016). Indeed, mutation of serine 131 to alanine on LSD1 inhibited B56a binding (Fig. S1C, D). It has been established that many chromatin-associated proteins are constitutively phosphorylated due to nuclear PP2A inhibition (Kauko et al., 2020; Zheng et al., 2015). PP2A/B56 complex is inhibited in RAS mutant lung cancer cells by direct interaction with SET (Saddoughi et al., 2013; Vainonen et al., 2021), and based on ours, and published results, SET and B56 are found from the nucleus (Xu et al., 2024)(Fig. S1E). By proximity labelling assay (PLA) we further found that SET associates with LSD1 in the nucleus (Fig. S1F). To validate phosphorylation regulation of S166 on LSD1 by SET-mediated PP2A inhibition, our attempts to develop a S166 specific phosphoantibody repeatedly failed. However, we could confirm S166 dephosphorylation in two SET depleted cell lines by targeted proteomics (SRM) assay developed for this purpose (Fig. S1G). These results identify S166 of LSD1 as a novel target site for SET-regulated PP2A/B56 in cancer cells.

### PP2A activation and RAS inhibition increase chromatin recruitment of LSD1

To study the functional consequences of PP2A/B56 on LSD1 in KRAS mutant cancer cells H460, we either depleted SET by siRNA, or treated cells with FTY720 which is an established inhibitor of SET-PP2A/B56 interaction (Saddoughi et al., 2013; Vainonen et al., 2021). Interestingly, while SET inhibition did not impact LSD1 protein expression (Fig. S1H, I), it resulted in robust increase in chromatin recruitment of LSD1 by using cell fractionation assay (Fig. 1D-G). Increased LSD1 chromatin recruitment was observed also using other PP2A/B56 activating compounds DT061or DBK1154, that function as molecular glues increasing B56 affinity to PP2A core dimer (Fig. 1F,G)(Denisova et al., 2023; Vainonen et al., 2021). The impact of PP2A activation in LSD1 chromatin recruitment was validated across two other KRAS mutant lung cancer cell lines (Fig. 1H, I). Moreover, inhibition of RAS by siRNA increased chromatin recruitment of LSD1 (Fig. S1J, K). Conversely, overexpression of oncogenic KRAS G12V mutant, and inhibition of PP2A/B56 by small-t antigen (Hahn et al., 2002), resulted in stepwise decrease in chromatin recruitment of LSD1 in human bronchial epithelial cells (Fig. 1J,K).

To demonstrate whether S166 dephosphorylation alone is sufficient to induce chromatin recruitment of LSD1, the H460 cells were transiently transfected with either wild-type (WT), alanine (S166A), or glutamic acid (S166D) mutants of LSD1, mimicking dephosphorylated and constitutively S166 phosphorylated LSD1, respectively. Consistent with results in PP2A-activated and RAS-inhibited cells, the S166A mutation resulted in strong increase in chromatin retention of LSD1, whereas the S166D mutant resembled LSD1 WT nuclear partitioning (Fig. 1L,M).

These results show that PP2A/B56 and RAS activities co-operatively regulate the chromatin occupancy of LSD1.

### S166 dephosphorylation induces chromatin recruitment of LSD1 and HDAC1/2

To validate these results in the endogenous setting, we introduced S166A mutation to *LSD1* gene locus in KRAS mutant lung cancer cell line H460 by CRISPR/CAS9 (Fig. 2A). Indicative of essential function for S166 phosphorylation switch in these KRAS mutant cells, only heterozygous S166A mutant clones could be obtained as verified by targeted DNA sequencing. The functional heterozygosity was verified by approximately 50% reduction in S166 phosphorylation by targeted proteomics assay (Fig. 2B). Two independent single cell clones of heterozygous S166A mutant cells did not show any apparent phenotype in long term colony growth assay confirming that any subsequent functional phenotypes are not due to cellular toxicity (Fig. 2C).

**Figure 2.**
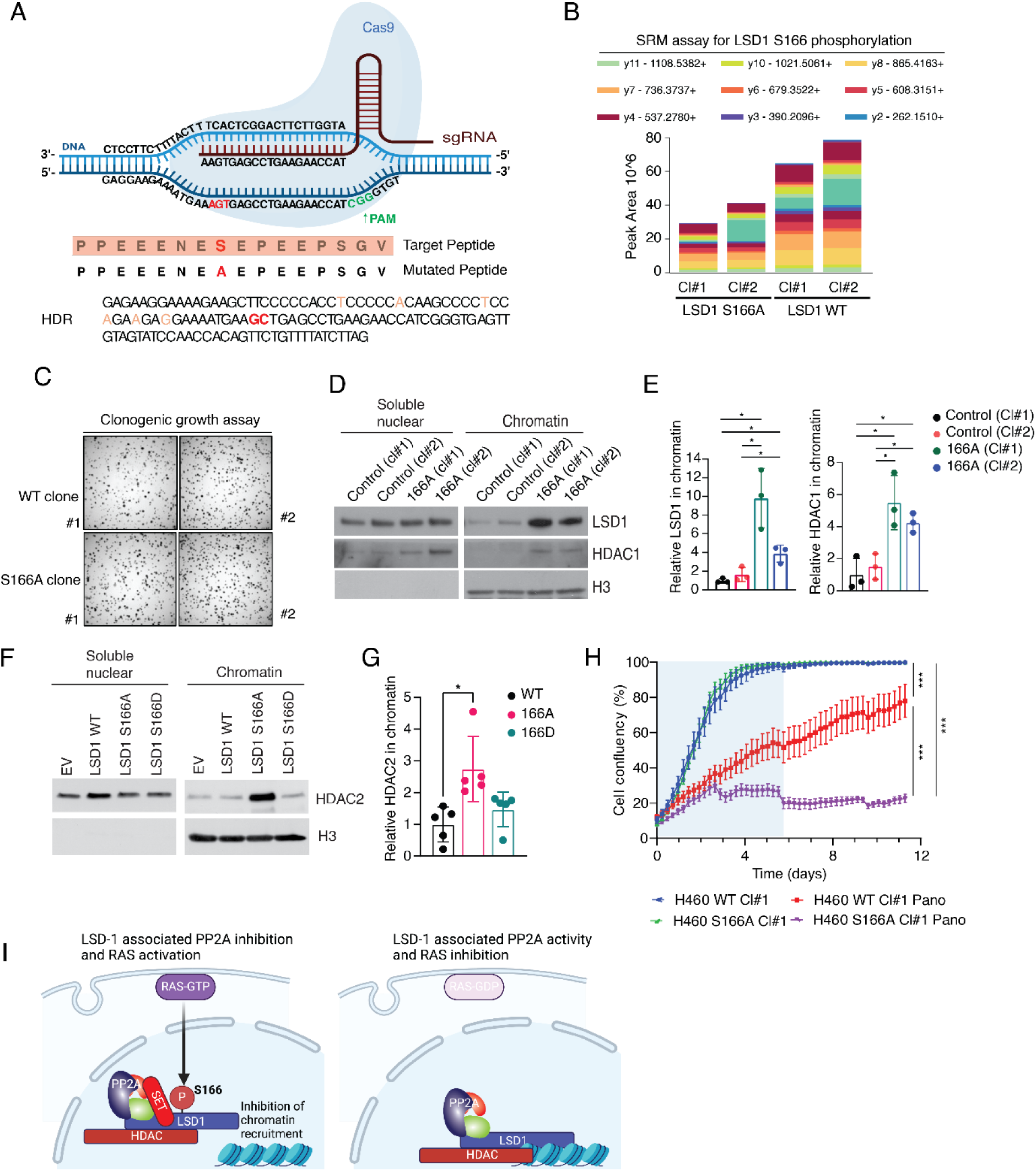
RAS and PP2A mediated phospho-regulation at S166 controls the chromatin recruitment of LSD1. **(A)** Schematic representation of the CRISPR/Cas9-based gene editing strategy used to generate LSD1 S166A mutant H460 cells. The sgRNA sequence targeting LSD1 is shown, along with the homology-directed repair (HDR) template introducing the S166A mutation. The target peptide sequence and corresponding mutated sequence are indicated. **(B)** Selected Reaction Monitoring (SRM) assay quantifying LSD1 S166 phosphorylation levels in WT and S166A mutant H460 clones. The colored bars represent phosphopeptide ion intensities for different transitions measured across clones. **(C)** Clonogenic growth assay comparing the colony-forming ability of WT and LSD1 S166A mutant clones. Cells were grown for 14 days, fixed, and stained to visualize colonies. **(D)** Western blot analysis of chromatin recruitment of endogenous LSD1 and HDAC1 in H460 WT or LSD1 S166A mutant clones. **(E)** Quantification of results from (D). (n = 3). **(F)** Western blot analysis of chromatin recruitment of endogenous HDAC2 in H460 cells, transiently transfected with either wild type or S166A/D mutants of LSD1. **(G)** Quantification of results from (F) (n = 5). **(H)** Incucyte analysis of the proliferation of H460 WT clone #1 and S166A clone #1 treated with DMSO and Panobinostat (50 nM) (12 wells/condition). The cells were treated with drugs twice per week followed by six days of drug holiday. Data were analyzed using one way ANOVA, and results are presented as mean ± standard deviation (SD). Statistical significance determined using the thresholds, *p < 0.05, **p < 0.01, ***p < 0.001 and P ≥ 0.05, not significant (ns). **(I)** Schematic representation describing how RAS and PP2A regulated LSD1 S166 phosphorylation impacts recruitment of LSD1 and HDAC to the chromatin. **(E, G)** Statistical Analysis: Data were analyzed using Student’s t-test, and results are presented as mean ± standard deviation (SD). Statistical significance was determined using the following thresholds: *p < 0.05, **p < 0.01, and ***p < 0.001.

Consistent with transient ectopic expression data (Fig. 1L,M), both mutant H460 single-cell clones displayed increased chromatin recruitment of S166A mutant LSD1 protein (Fig. 2D,E). Further, the cells with endogenous S166A mutation also displayed co-recruitment of central NuRD complex component HDAC1 to the chromatin (Fig. 2D,E). Importantly, also transient overexpression of S166A mutant LSD1 resulted in increased chromatin recruitment of HDAC, whereas this was not seen with S166D mutant (Fig. 2F,G). These results demonstrate that lack of S166 phosphorylation on LSD1 has dominant effect at the protein complex level, and thereby potentially wider impact of cancer cell epigenome regulation. As the functional consequence of 50% reduction in LSD1 S166 phosphorylation, and HDAC protein recruitment to the chromatin, the LSD1 S166A mutant cells were significantly more sensitive to HDAC inhibitor panobinostat than control cells (Fig. 2H). Lower panobinostat sensitivity of the wildtype cells in which HDAC is not enriched on the chromatin due to S166 phosphorylation, is fully consistent with previously reported HDAC inhibitor resistance induced by PP2A inhibition (Kauko et al., 2020).

Collectively these results show that S166 of LSD1, regulated by RAS and SET-mediated PP2A/B56 inhibition, is a critical determinant of chromatin occupancy of both LSD1 and HDAC (Fig. 2I).

### LSD1 S166 phosphorylation controls epigenetic landscape

LSD1 chromatin immunoprecipitation sequencing (ChIP-seq) analysis using two independent single cell clones of LSD1 S166A mutant and wildtype H460 cells showed 1270 peaks/genomic regions which show differential LSD1 binding (Fig. 3A,B and S2A). Consistent with the increased recruitment observed by chromatin fractionation, differential binding analysis on the consensus peaks set revealed that approximately two thirds of the differentially occupied sites showed increased LSD1 S166A mutant binding (1029 regions) (Fig. 3A,B and S2A). However, as overall there was a substantial conservation of bound regions between the wildtype and the mutant clones (Fig. S2A), we conclude that the S166A mutation could be increasing the overall chromatin association of the protein (evident in fractionation assays) by stabilizing or enhancing binding at already occupied sites, but also by recruiting the protein to new genomic locations. Examples of gene promoters with increased LSD1 S166A binding are shown in figure 3C. The genes in which S166A mutation increased chromatin binding were enriched in processes related to development, differentiation and actin cytoskeleton, but also to known LSD1 co-factors such as SNAI1 and HDAC (Fig. S2B).

**Figure 3.**
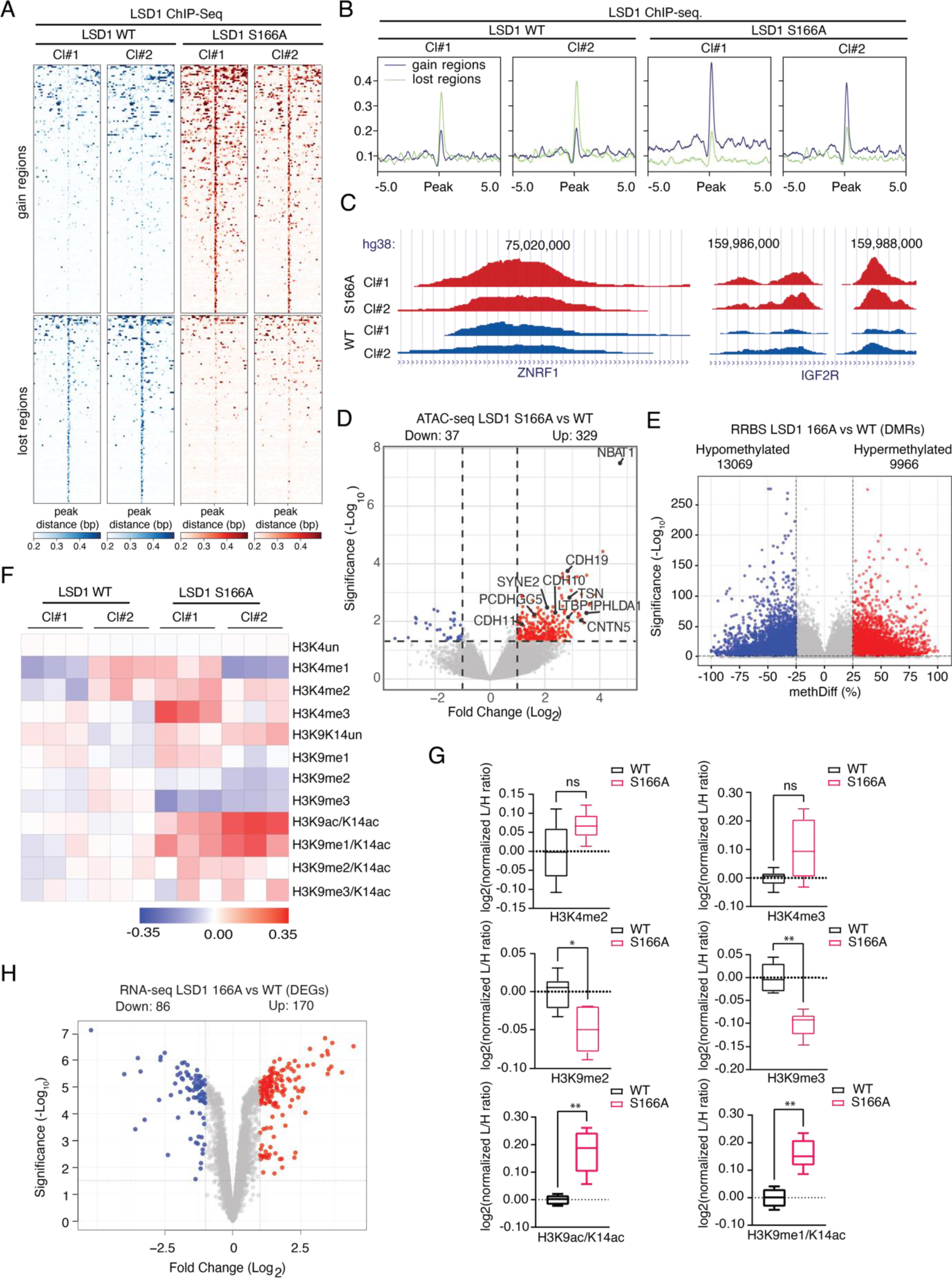
Impact of LSD1 S166 Dephosphorylation on Chromatin Landscape. **(A)** Tornado plot and profile plot of LSD1 ChIP-Seq on regions identified in Fig. 3B as gained (n = 165, purple lines in density plot, upper part of the heatmap) or lost (n = 125, green lines in density plot, bottom part of the heatmap) in the control (blue) and LSD1 mutant cell clones (red). The values were centered and plotted in a ±5 kb genomic interval relative to the summit of the peaks. **(B)** Profile plot of LSD1 ChIP-Seq signal on regions identified in fig A as gained (n= 165, purple) or lost (n=125, green) in the WT and LSD1 mutant cell clones. **(C)** Chip-seq tracks of two example genomic regions in which LSD1 S166A mutation increased chromatin occupancy. **(D)** Volcano plot of ATAC-seq analysis comparing chromatin accessibility between LSD1 S166A and WT cells (n = 3). Genes with significantly increased chromatin accessibility are in red, while those with decreased accessibility are in blue (log2FC > ±1, p < 0.05). The most significantly accessible genes and those involved in extracellular matrix (ECM) organization are labeled. Differentially accessible regions were determined using DESeq2 version 1.40.2. **(E)** Volcano plot of reduced representation bisulfite sequencing (RRBS) analysis, showing differentially methylated regions (DMRs) between LSD1 S166A and WT cells (Methylation difference (MethylDiff) > ±25%, FDR < 0.05) (n = 3). Differential methylation analysis was run using Bioconductor R-package methylKit. **(F)** Heatmap showing the relative abundance of histone 3 (H3) post-translational modifications in LSD1 S166A mutant and WT clones based on mass spectrometry analysis of histones. **(G)** Quantification of H3 marks from G. The data from the WT and S166A mutant clones were combined to increase the statistical power of the analysis (n=6). Data were analyzed using Mann–Whitney test, and results are presented as mean ± standard deviation (SD). Statistical significance was determined using the following thresholds: *p < 0.05, and **p < 0.01. **(H)** Volcano plot showing differentially expressed genes (DEGs) between LSD1 S166A mutant and WT cells, based on RNA-seq data (n = 3). Genes with significant upregulation are in red, while downregulated genes are in blue (log2FC > ±1 & FDR < 0.05). Statistical testing was performed using the ROTS package.

To further explore epigenetic impact by LSD1 S166 dephosphorylation, we performed Assay for Transposase-Accessible Chromatin using sequencing (ATAC-seq) analysis. Importantly, the analysis revealed selective, consistent changes, and an increased chromatin accessibility at 329 regions in the S166A mutant cells (Fig. 3D and Fig. S2C). The general distribution of the differentially accessible peaks (DAPs) between control and LSD1 S166A mutant cells across genomic regions and chromosomes resembled that of ATAC-seq profiles in general (Fig. S2D,E), and the open chromatin regions in S166A LSD1 mutant cells were enriched for genes functionally involved in cell junction assembly and Cadherin signaling (Fig. S3A, B). In addition to analyzing the most accessible chromatin regions, we performed a footprinting analysis on ATAC-seq data to further explore protein-DNA interactions. The results revealed a substantial number of transcription factors predicted to exhibit increased binding in the LSD1 S166A mutant compared to the WT counterpart (Fig. S3C). Notably, transcription factors showing greater binding in the LSD1 S166A mutant are associated with positive regulation of transcription, indicating a potential shift toward a transcriptionally more active chromatin state (Fig. S3D).

In addition to its direct function as a H3 demethylase, LSD1 impacts gene expression also by indirect mechanisms, such as by maintaining DNA methylation (Perillo et al., 2020; Wang et al., 2009). Therefore, we analysed DNA methylation in LSD1 S166A mutant cells by reduced-representation bisulfite sequencing (RRBS). By this assay, phosphorylation status of S166 of LSD1 appears to have dramatic effects on epigenome. As compared to WT cells, LSD1 S166A mutant clones demonstrated hypomethylation of over 13 000 genomic regions and hypermethylation of almost 10 000 genomic regions (Fig. 3E and S4A). Among the hypomethylated regions, genes related to ECM biology were highly enriched (Fig. S4B).

To eventually study the impact of S166A mutation on LSD1-mediated histone modifications, we analyzed H3 histone marks by Super-SILAC mass spectrometry approach for profiling histone posttranslational modifications (Noberini et al., 2023). Interestingly, the *bona fide* target of LSD1 catalytic activity, H3K4 methylation, was not significantly changed between mutant and control cells, but there was significant inhibition of H3K9 di -and trimethylation in LSD1 S166A mutant cells. This was associated with significantly increased H3K9/14 acetylation (Fig. 3F,G).

All the above analyzed epigenetic effects by LSD1 S166A mutation are indicative of increased gene activation. To study how S166 dephosphorylation eventually affects LSD1 function in gene regulation, we performed RNA sequencing (RNAseq) analysis comparing two control and two S166A mutant clones all in triplicate. When overlapping significantly regulated (Log2 0,5; FDR 0.05) genes from the control clones and mutant clones where compared, LSD1 S166A mutation was found to significantly alter the expression of the 824 genes (differentially expressed genes; DEGs) (Fig. 3H, Table S1). From them, 536 DEGs were upregulated, and 288 genes downregulated in S166A mutant clones, indicating that S166 dephosphorylation unleashes the gene activator function of LSD1. This conclusion was enforced by Gene Set Correlation Enrichment analysis (Chang et al., 2024) of the DEGs, indicating that the gene expression profile in S166A cells significantly resembled the situation in which the normal gene repression function of LSD1 was inhibited (Fig. S4C). Consistent with gene activation data from LSD1 S166A mutant cells mimicking PP2A mediated dephosphorylation of S166, PP2A reactivation by SET inhibition in HeLa cells also resulted primarily in gene activation (Fig. S4D, Table S2). Although the RNA sequencing data of these two PP2A activation models were obtained from different cell lines (S166A, H460; siSET, Hela), the overlap of genes significantly overexpressed was notable (17% of the siSET regulated genes) (Fig. S4E), whereas there was no overlap between the datasets regarding the downregulated genes. Differential expression of 28 genes was further validated by qPCR from independent triplicate WT and S166A samples (Fig. S4F, G). Importantly, six of these genes were found significantly upregulated by SET siRNA in HeLa cells demonstrating their positive regulation by both S166 dephosphorylation and PP2A activation (Fig. S4H).

These results demonstrate general epigenetic landscape effects by LSD1 S166 dephosphorylation, and that S166 dephosphorylation functions as a phosphorylation switch for gene activator function of LSD1.

### Inhibition of protein interactions with transcriptional repressors SNAI2 and MYBPP1 by S166A mutation of LSD1

We next aimed to understand how S166A dephosphorylation might change LSD1 function in chromatin remodeling, histone modifications, and gene expression. As expected from the other studies targeting the amino terminus of LSD1 (Waterbury et al., 2024), and lack of effects on H3K4 methylation (Fig. 3F,G), S166A mutation did not impact the catalytic amine oxidase activity of recombinant LSD1 protein (Fig. 4A). S166A mutation also did not impact the sensitivity of LSD1 to catalytic LSD1 inhibitor molecule GSK-LSD (Fig. 4B). We also did not observe any indications for major structural effects of S166A mutation on LSD1 based on differential scanning fluorometer (DSF) analysis (Fig. 4C). These results strongly indicate that S166A mutation do not affect histone code and gene regulation by impacting the catalytic activity of LSD1.

**Figure 4.**
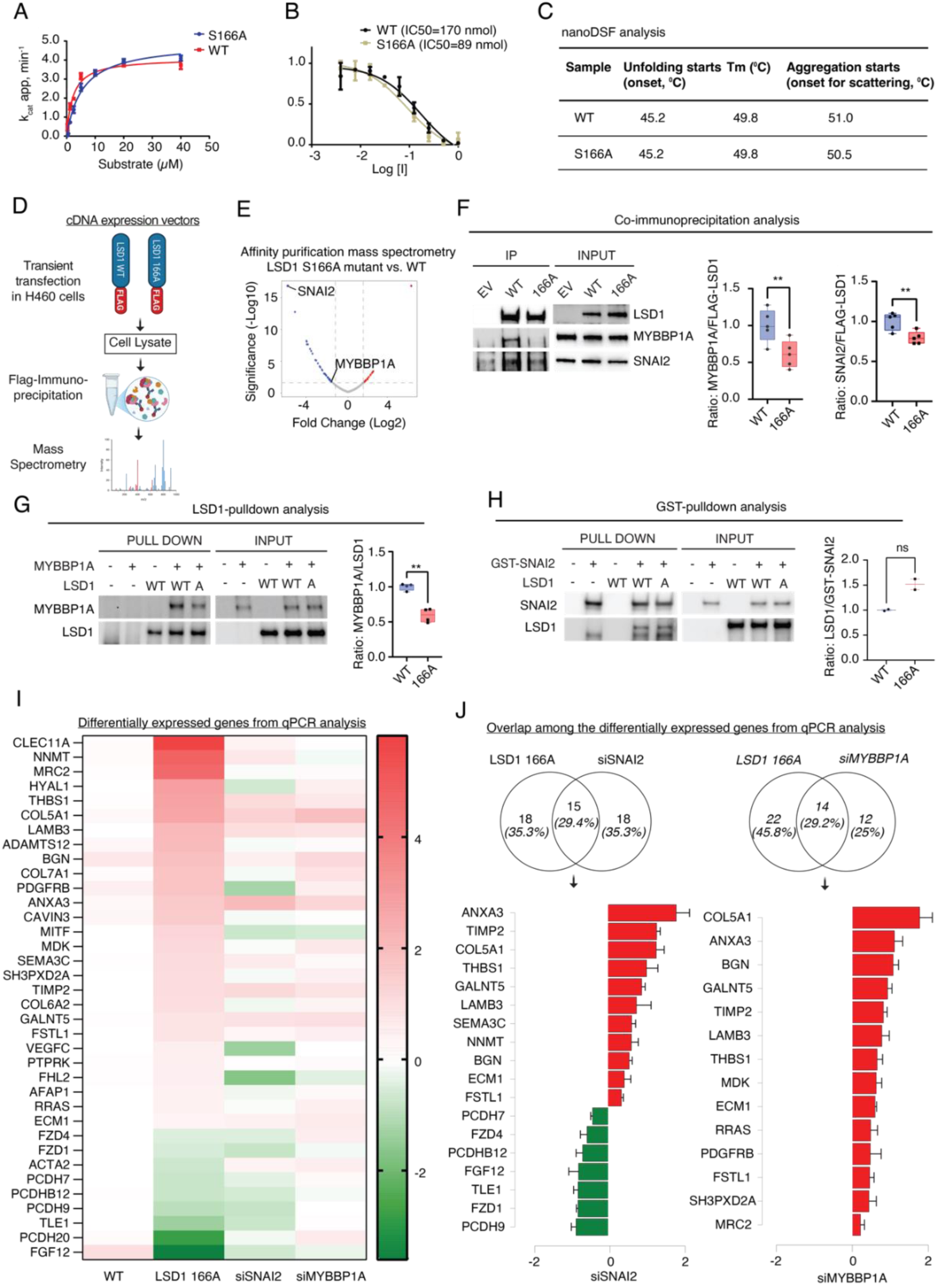
LSD1 S166 dephosphorylation impairs its interaction with transcriptional repressors SNAI2 and MYBBP1A. **(A)** Catalytic rate of purified wild-type (WT) and mutant (S166A) LSD1 protein incubated with H3K4me2 as substrate. **(B)** Catalytic rate of purified wild-type (WT) and mutant (S166A) LSD1 protein incubated with H3K4me2 as substrate treated with GSK-LSD1 inhibitor. **(C)** Nano differential scanning fluorometry (nDSF) analysis of temperature sensitive protein folding of the purified wild-type (WT) and mutant (S166A) LSD1 protein. **(D)** Schematic representation of Immunoprecipitation mass spectrometry analysis (AP-MS) of the interaction partners for LSD1 wild-type (WT) and mutant (S166A) transiently transfected in H460 cells. **(E)** Volcano plot showing the differential interactome between S166A mutant and wild-type (WT) LSD1 determined by AP-MS as described in (D). **(F)** Co-immunoprecipitation analysis of the impact of transiently transfected H460 cells with flag tagged LSD1 wild-type (WT) or mutant (S166A) on protein interaction with MYBBP1A and SNAI2. The strength of interaction was quantified as the ratio between MYBBP1A/SNAI2 and flag tagged LSD1 WT or mutant (n=5). Data were analyzed using Student’s t-test, and results are presented as mean ± standard deviation (SD). Statistical significance was determined using the following threshold: **p < 0.01. **(G)** In vitro pull-down assay between purified MYBBP1A and LSD1 wild-type (WT) or mutant (S166A) proteins using LSD1 antibody. The strength of interaction was quantified as the ratio between MYBBP1A and LSD1 WT or mutant proteins (n = 4). Data was analyzed using Student’s t-test, and results are presented as mean ± standard deviation (SD). Statistical significance was determined using the following threshold: **p < 0.01. **(H)** In vitro pull-down assay between purified GST tagged SNAI2 and LSD1 wild-type (WT) or mutant (S166A) proteins using GST trap agarose (n = 2). The strength of interaction was quantified as the ratio between LSD1 WT or mutant and GST-SNAI2 (n = 2). Data was analyzed using Student’s t-test, and results are presented as mean ± standard deviation (SD). Statistical significance was determined using the following threshold: P ≥ 0.05, not significant (ns). **(I)** Heatmap describing the change in expression of genes in LSD1 S166A mutant H460 cells, SNAI2 silenced H460 cells and MYBBP1A silenced H460 cells and WT H460 as determined by Q-PCR analysis. **(J)** Venn diagram comparing the overlap between the differentially expressed genes in S166A mutant H460 cell clones and SNAI2 or MYBBP1A silenced H460 cells. The bar graph below the venn diagrams shows the changes in expression of the common genes between LSD1 S166A mutant H460 cell clones and SNAI2 or MYBBP1A silenced H460 cells (n = 3). Data shows significantly regulated genes analyzed using Student’s t-test, and results are presented as mean ± standard deviation. A value of p < 0.05 was considered as significant.

One of the important functions of protein phosphorylation is the regulation of protein interactions (Hunter, 2012; Westermarck et al., 2013), and it is well established that LSD1 functions are strongly modulated by protein interactions. Thereby, we assessed whether S166A mutation might affect protein interactions between LSD1 and its cognate partners in chromatin complexes. To this end, H460 cells were transiently transfected with either Flag-tagged WT or S166A mutant LSD1 and protein interactions from chromatin fractions were studied by affinity-purification coupled with mass spectrometry (AP-MS) analysis (Fig. 4D).

Importantly, scaffolding A-subunit of PP2A was found from WT LSD1 interactome but it interacted less with the S166A mutant LSD1 (Fig. S5A), re-enforcing the role for PP2A as the S166 phosphatase. S166A mutation had a clear impact on LSD1 protein interactome characterized by loss of protein interactions (Fig. 4E, Table S3). Interestingly, the proteins that had impaired interaction with S166A mutant LSD1 as compared to WT LSD1 were enriched in processes of Cadherin binding and Cell adhesion molecule binding (Fig. S5B). In search for altered protein interactions underpinning the profound epigenetic effects of S166 dephosphorylation of LSD1, we noted significantly impaired interaction of LSD1 S166A with two epigenetic repressor proteins MYBBP1A and SNAI2 (Fig. 4E). SNAI2 is a known gene repressor partner for LSD1 (Ferrari-Amorotti et al., 2013), whereas MYBBP1A increases DNA methylation (Weng et al., 2019), and regulates histone code (Tan et al., 2012; Yang et al., 2012), consistent with phenotypes seen in S166A cells (Fig. 3). Thereby, loss of these two repressor proteins from the LSD1 complex could explain most of the observed epigenetic changes in S166A mutant cells.

Co-immunoprecipitation analysis of transiently transfected FLAG tagged WT or S166A mutant LSD1 and using MYBPP1A and SNAI2 antibodies for western blot detection, confirmed significantly reduced interaction of these repressor proteins with S166A mutant LSD1, as compared to WT protein (Fig. 4F). Importantly, consistently with proposed role for the disordered N-terminal region regulating LSD1 interactions with selected co-factors, but not with the core CoREST/NuRD repressor components (Waterbury et al., 2024), S166A mutation did not impact protein interaction between LSD1 and HDACs based on AP-MS data (Table S3).

We further purified recombinant proteins and validated direct protein interaction between LSD1 and both MYBBP1A and SNAI2 (Fig. 4G,H). Direct interaction with SNAI2 was not impacted by S166A mutation of LSD1, whereas the mutation significantly decreased direct interaction between MYBBP1A and LSD1 (Fig. 4G,H). This indicates that while SNAI2 directly interacts with LSD1, yet unidentified proteins decrease its affinity to LSD1 S166A *in cellulo*. On the other hand, selective *in vitro* impact of S166A mutation on MYBBP1A, but not on SNAI2, further indicates that S166A mutation does not have overall disruptive effects on LSD1 protein.

To test whether loss of SNAI2 and MYBPP1 interactions could explain the gene expression phenotype in LSD1 S166A cells, we studied the impact of siRNA-mediated SNAI2 and MYBPP1A depletion on the panel of mutant LSD1 sensitive genes selected from the RNAseq data. By averaging three biological replicates from both S166A mutant clones (n=6), as well as SNAI2 and MYBPP1A depleted cells (n=3 for both), there was a very clear overlap between genes regulated by either S166A mutation or by loss of these gene repressor proteins (Fig. 4I,J). Whereas SNAI2 depletion mimicked the impact of S166A mutation for both up - and downregulated genes (overlap 18/36 genes), MYBBP1 depletion overlapped with the mutation only regarding the upregulated genes (overlap 14/36) (Fig. 4J). Collectively 24/36 of the studied genes could be attributed to either SNAI2 or MYBBP1A function (Fig. 4J) demonstrating their pivotal roles in the altered transcriptional activity of LSD1 S166A mutant protein.

These results uncover the mechanistic basis how dephosphorylation of S166 unleashes the transcriptional activator function of LSD1 by inhibition of protein interactions with gene repressors SNAI2 and MYBBP1A.

### Dephosphorylation of S166 on LSD1 results in activation of gene expression program related to ECM biology

Interestingly, the pathway enrichment analyses based on DEGs revealed a clear selectivity in gene expression programs regulated by LSD1 phosphorylation. Consistently with loss of interactions with proteins implicated in cell adhesions (Fig. S5B), the genes upregulated by S166A mutation were enriched in related processes such as extracellular matrix (ECM), and integrin biology (Fig 5A,B and Fig. S6A). Further emphasizing regulation of these biological functions by LSD1 S166 phosphorylation, they were found to overlap on pathway clustering and Metascape analysis based on RNAseq, ATAC-seq, and RRBS data (Fig. S6B,C). Interestingly, based on both Reactome and iPathway analysis, S166A mutant cells displayed also differential expression of genes related to immune and cytokine signaling (Fig. 5A,B and S6A). These results indicate that rather than affecting global gene regulatory mechanisms, S166A mutation may activate gene expression in a more selective manner. To study this, we performed a transcription factor binding site enrichment analysis (TFEA) using genes significantly upregulated in S166A mutant cells (Fig. 5C). Importantly, among the highly enriched transcription factors, SNAI2 and Notch-linked HEYL and HES4, associate functionally with the ECM phenotypes (Herrera et al., 2016; Karsan, 2008)(Fig. 5C). Further, enrichment of SNAI2 targets among the upregulated genes is fully consistent with impaired SNAI2 interaction with the LSD1 S166A mutant (Fig. 4), whereas enrichment of FOXO family binding sites is consistent with ATAC-seq. footprinting analysis (Fig. S3C).

**Figure 5.**
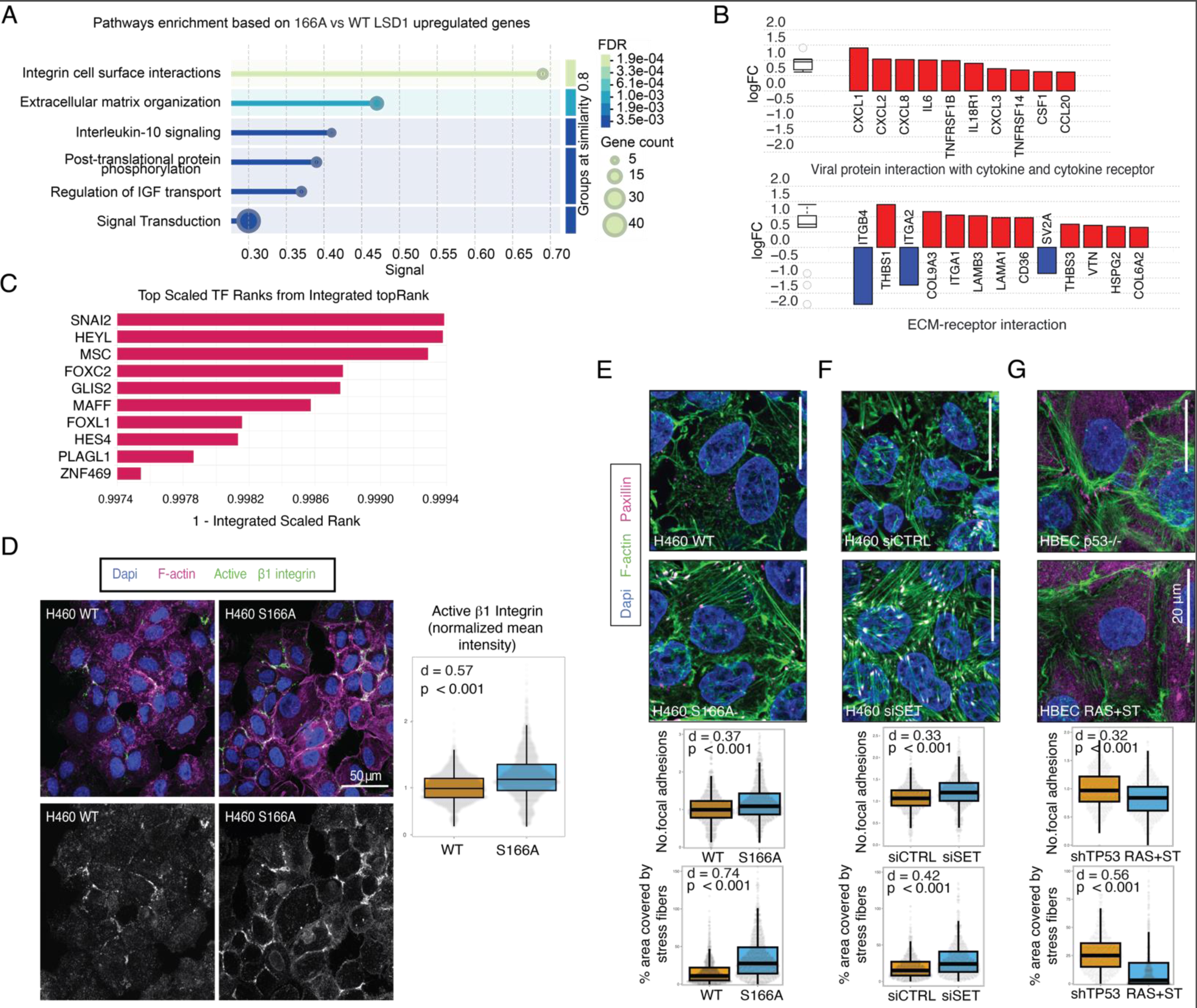
LSD1 S166 dephosphorylation induces gene activation related to ECM organization and cell contractility. **(A)** Pathway enrichment analyses based on upregulated genes from LSD1 S166A vs WT cells from RNA sequencing data. **(B)** Functional analysis of the RNA sequencing data of S166A vs WT H460 cells using iPathwayGuideTM (AdvaitaBio Corporation) highlighting the log2 fold-change (log2FC) values of genes involved in viral protein interactions with cytokine and cytokine receptors (top panel) and ECM-receptor interactions (bottom panel). Genes with increased expression are in red, while those with decreased expression are in blue. **(C)** Bar chart corresponding to the top ranked transcription factors identified in the LSD1 S166A vs. WT RNA sequencing dataset. **(D)** Immunofluorescence staining of active beta1integrin in H460 WT and LSD1 S166A mutant cells. **(E)** Immunofluorescence staining of F-actin (stress fibers) and paxillin (focal adhesions) from H460 WT and LSD1 S166A mutant cells. (**F**) Increased stress fibers and focal adhesions in response to PP2A reactivation by SET inhibition. (**G**) Decreased stress fibers and focal adhesions in response to mutant RAS overexpression and PP2A inhibition by small-t overexpression. **(D-G)** Cells were imaged using a spinning disk confocal microscope. Randomization tests and Cohen’s D values were computed using code available at (https://github.com/CellMigrationLab/Plot-Stats).

Results above demonstrate that dephosphorylation of S166 on LSD1 is sufficient alone to alter the balance between gene repressor and activator functions of LSD1, and to result in activation of selective gene expression program related to ECM biology. Interestingly, one of the upregulated genes was β1 integrin activator RRAS (Liu et al., 2017) (both in RNAseq and qPCR data (Table S1, and Fig. S4F, G). Indeed, by using immunofluorescence staining with an antibody recognizing specifically the activated conformation of β1 integrin, we detected significantly more activation in S166A than in WT H460 cells (Fig. 5D). Furthemore, increased integrin activity translated to significantly more prominent paxillin-positive focal adhesions and more actin stress fibers (Conway and Jacquemet, 2019)(Fig. 5E). To link these findings back to PP2A -and RAS-mediated S166 regulation, both phenotypes were activated by PP2A reactivation in SET-depleted cells (Fig. 5F), whereas they were suppressed in HBEC cells transduced with mutant HRAS and PP2A inhibitor small-t (Fig. 5G).

In summary, lack of LSD1 S166 phosphorylation induces transcriptional activation of genes that manifest in integrin activation and cell contractility.

### LSD1 S166 dephosphorylation in cancer cells regulates tumor microenvironment and macrophage recruitment

As epigenetic alterations in cancer cells have been shown to impact tumor microenvironment and tumor immune cell composition (Yang et al., 2023), we proceeded to study the *in vivo* relevance of LSD1 S166 dephosphorylation in cancer cells. To this end, we performed subcutaneous xenograft experiments with the WT and S166A LSD1 mutant H460 cells in nude mice. As *in vitro* (Fig. 2C), lack of LSD1 S166 phosphorylation did not have detrimental effects on xenograft growth allowing studying the associated gene expressions changes also *in vivo.* In fact, The LSD1 S166A mutant cells displayed repeatedly (3 out of 3 experiments) faster tumor growth than WT tumors, but this difference did not reach the statistical significance (Fig. 6A and Fig. S7A). Whether this observation has cancer relevance remains a subject for future studies, but interestingly significantly lower S166 phosphorylation levels were found in stage 3 lung adenocarcinomas based on Clinical Proteomic Tumor Analysis Consortium (CPTAC) data (Ellis et al., 2013)(Fig. S7B), and in breast tumors compared to normal mammary tissue (Fig. 1B). Importantly, using our qPCR target panel on mRNA samples from five parallel tumor samples, the gene activation function of LSD1 S166A mutation was very apparent also *in vivo* (Fig. 6B and S7C).

**Figure 6.**
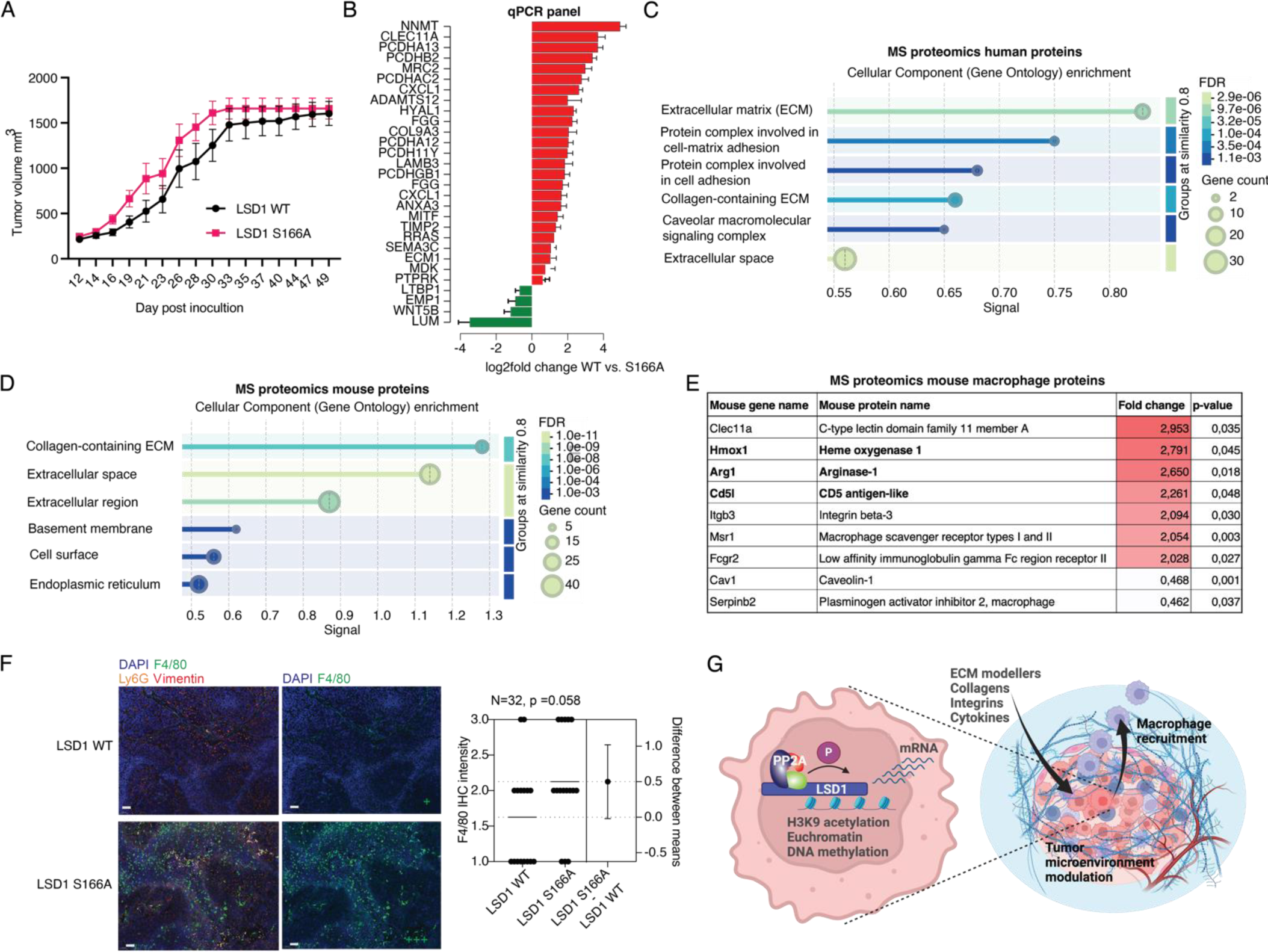
LSD1 S166 dephosphorylation shapes the tumor microenvironment and enhances macrophage recruitment. **(A)** Tumor growth follow-up of subcutaneous xenografts of H460 WT clone #1 (n=7) or S166A clone #1 (n=8). **(B)** Gene expression analysis between LSD1 S166A mutant and WT xenografts from (A) (n = 5 each genotype) as determined by Q-PCR. Data shows significantly regulated genes analyzed using Student’s t-test, and results are presented as mean ± standard deviation. (SD). A value of p < 0.05 was considered as significant. **(C)** Cellular component enrichment analysis of human proteins based on mass spectrometry analysis of S166A vs WT LSD1 xenografts from (A) (n = 5 each genotype). **(D)** Cellular component enrichment analysis of mouse proteins based on mass spectrometry analysis of S166A vs WT LSD1 xenografts from (A) (n = 5 each genotype). **(E)** Expression changes of significantly altered mouse macrophage markers based on mass spectrometry analysis of S166A vs WT LSD1 xenografts from (A) (n = 5 each genotype). In bold M2 type selective mouse macrophage markers. **(F)** Multiplex IHC staining of tumor sections from (A) with macrophage marker F4/80, neutrophil marker Ly6G and DAPI. Shown is representative samples of low (+) F4/80 positivity in LSD1 WT tumor and high (+++) positivity in LSD1 S166A tumor. Scale bar 500 µm. Quantification is based on visual inspection of multiple independent stainings combining tumors from both independent WT and S166A mutant clones. **(G)** Schematic presentation of paracrine regulation of tumor stroma and macrophage infiltration by S166 dephosphorylation of LSD1via epigenetic regulation of ECM genes.

Whereas gene expression data was performed using a preselected qPCR gene panel, we further performed MS proteomics analysis of the formalin-fixed tumor tissues to assess whether the LSD1 S166A mutation-elicited epigenetic effects impact tumor phenotype at the proteome level. Using three replicate tumors per genotype, 27 proteins were found to be upregulated and 36 proteins downregulated in the tumor tissues expressing LSD1 S166A (Table S4). Notably, like gene expression changes, also proteins differentially regulated by S166A were highly enriched in extracellular matrix, cell adhesion and collagen regulation (Fig. 6C).

To evaluate whether epigenetic regulation by LSD1 S166A mutant also can influence the host tissue, we surveyed changes in mouse proteome expression in tumor tissues by annotation of peptides against mouse proteome reference database. Intriguingly, indicating for a robust capability of LSD1 S166A mutant human cancer cells to induce remodelling of the surrounding mouse stroma, altogether 99 mouse proteins were found more abundantly present in the S166A mutant tumors as compared to WT tumors (Table S5). Consistent with altered tumor stroma phenotype in S166A tumors, the differentially expressed proteins were highly enriched in ECM proteins (Fig. 6D).

Altered tumor stroma and ECM are known to affect immune cell tumor infiltration (Deligne and Midwood, 2021; Mai et al., 2024). This was enforced by upregulation in S166A LSD1 mutant cancer cells of many genes in involved in macrophage infiltration such as *CXCL1, MRC2, SEMA3C, TIMP2, ECM1, MDK,* and *LUM* (Fig. 6B). At human proteome level, we identified several upregulated proteins that are involved in macrophage infiltration and activation. These include for example MRC1, IL-18, SERPINB2, TGM2, and PLAU (Table S4). This prompted us to survey the mouse proteome data for potential enrichment of mouse macrophage specific proteins as a quantitative measure of macrophage tumor infiltration. Indeed, we identified significant increase in abundance of many *bona fide* mouse macrophage proteins in the S166A tumors as compared to WT tumors. Further, significant upregulation of Hmox1, Arg1, and Cd5l indicated that in addition to increased infiltration, the macrophages in S166A mutant tumors may predominantly be M2 type macrophages. This is consistent with role of increased tissue stiffness in promoting M2 macrophage polarization (Mai et al., 2024), and our data demonstrating a more actin rich and contractile-like phenotype in LSD1 S166A cells (Fig. 5D-G). Further, IL-10 signaling was one the enriched pathways based on genes upregulated in S166A mutant cancer cells (Fig. 5A), and IL-10 is known to promote M2 polarization (Wang et al., 2021). This quantitative proteomics data was validated by multiplex IHC staining of tumor sections with macrophage marker F4/80, neutrophil marker Ly6G, and DAPI (Fig. 6F, S7D). The results validated increased mouse macrophage recruitment in LSD1 S166A mutant tumors, whereas suggestive of selective effects on macrophages, we did not observe obvious difference between tumors regarding their neutrophil content (Fig. 6F, S7D).

As a summary, our data reveal that a single phosphorylation site in a non-histone protein in cancer cells can control tumor stroma content and immune cell infiltration via epigenetic gene activation (Fig. 6G).

## Discussion

Our results identify S166 of LSD1 as a central regulatory hub for epigenetic landscape regulation and tumor tissue organization. While genetic mutation landscapes in cancer and many other diseases have been studied for decades, and especially in cancer we have already almost saturated knowledge on recurrent gene mutations (Martincorena et al., 2017), our knowledge on physiological and pathological impact of PTMs lags very significantly behind (Hornbeck et al., 2019). One of the conceptual questions related to protein phosphorylation is whether it is merely used to create charged amino acid platforms in proteins impacting their folding and interactions, or how often phosphorylation of a single amino acid is sufficient to regulate an important cellular function or tissue homeostasis (Bludau and Aebersold, 2020; Hunter, 2012). Thereby, demonstration that a single phosphorylation site on unstructured and functionally uncharacterized region of a non-histone protein governs cancer cell epigenetic landscape, and tumor microenvironment, serves as an intriguing example of the power of individual PTMs in human biology.

Our results provide mechanistic explanation for longstanding enigma how LSD1 can function as either gene repressor or gene activator in context dependent manner in non-hormonal cancers. The results establish dephosphorylation of S166 as a molecular switch for unleashing the gene activator function for LSD1 via markedly increased chromatin accessibility, DNA hypomethylation, and H3K9 acetylation. Consistently with previous studies related to functional role of LSD1 N-terminus (Astro et al., 2022; Wang et al., 2009; Waterbury et al., 2024), we found that dephosphorylation of S166 neither affected the catalytic activity of LSD1, nor its overall structure, but rather had a dramatic impact on its protein interaction network. In particular, we found that dephosphorylation of S166 reduced the interaction of LSD1 with transcriptional repressors SNAI2 and MYBBP1A. This is consistent with recently identified role for the unstructed LSD1 N-terminus in inhibiting TF interactions by shielding the structured LSD1 domains via intramolecular LSD1 domain interactions (Waterbury et al., 2024). Interestingly as S166 is the sixth amino acid of the unstructured N-terminus counting from the structurally resolved first amino acid S172, we speculate that it may serve as a phosphorylation-regulated hinge that impacts positioning and the electrostatic interactions of the N-terminus and thereby TF and co-repressor interactions (Fig. S7F). Collectively the results enforce the unprecedented importance of the N-terminus in LSD1 function, and S166 as a regulatory switch for LSD1 protein interactions.

While previous data revealed that a close homologue of SNAI2, SNAI1, interacts with the amine oxidase domain of LSD1 (Lin et al., 2010), we hereby provide first evidence about direct interaction between LSD1 and SNAI2, or MYBBP1A. Our results demonstrating upregulation of ECM-related SNAI2 target genes by LSD1 S166A mutation imparing SNAI2 interaction, are supported by role for LSD1 in regulation SNAI2 target genes (Egolf et al., 2019), and that peptide inhibition of SNAI2 interaction with LSD1 reduced epithelial mesenchymal transition and cell migration (Ferrari-Amorotti et al., 2013). In addition to SNAI2 and MYBBP1A, the impact of S166 dephosphorylation on epigenome could be mediated by increased recruitment of other transcription regulators to more accessible euchormatin as demonstrated by the ChIP-seq and ATAC-seq footprinting analysis (Fig. S3C). This conclusion is further enforced by the oncogenic gene signature analysis of differentially expressed genes in LSD1 S166A mutant cells revealing strong enrichment of genes known to be regulated by epigenetic repressor protein BMI-1 (Fig S7E).

Our results reveal an intriguing new mechanism how extracellular matrix and tumor microenvironment can be regulated in a paracrine manner by PTMs in cancer cells. The tumors with the LSD1 S166A mutant lung cancer cells displayed robust changes in extracellular matrix composition derived from both human and mouse cells. The human cell derived ECM changes are consistent with the function of LSD1 in regulating gene expression related to cell migration and invasion (Egolf et al., 2019). Furthermore, our in vivo studies identify LSD1 166 dephosphorylation-mediated regulation of macrophage tumor infiltration. These findings corroborate earlier research indicating that ECM remodeling influences the immune microenvironment. For instance, it has been shown that integrin activation drives matrix stiffness, leading to macrophage polarization (Deligne and Midwood, 2021; Mai et al., 2024). In addition, prominent activation of cytokine gene expression in LSD1 S166A mutant cells (Fig. 5B, 6B and S6A) most likely also contributes to the macrophage phenotype in tumors.

Existing phosphoproteome data clearly indicate that S166 is under dynamic regulation of S166 in response to different cellular states and external stimuli (Fig. 1B). On the other hand, this site is not mutated in cancer and based on our Crispr/Cas9 mutagenesis results, homozygous mutant cells were not viable. These data together indicate that this site is under continuous phosphorylation-dephosphorylation cycle and that dephosphorylation of even fraction of cellular LSD1 pool has dominant cellular functions. Therefore, a very imporant future goal is to better understand the mechanisms of LSD1 S166 phosphorylation regulation. As both PP2A and RAS are very central regulators of cellular phosphorylation-dependent signaling mechanisms (Fowle et al., 2019; Kauko et al., 2020; Kauko et al., 2015; Klomp et al., 2024b), identification of S166 as a novel converge point for their activities is itself a very strong clue for the biological importance of this phosphorylation site. On the other hand, S166 is just four residues before an exon-intron junction that is site of alternative splicing in LSD1 (Astro et al., 2022). Thereby, inclusion of exon 2a might in some developmental states be another type of mechanism that impact the biological activity of LSD1 by altering the phosphorylation-dephosphorylation balance on S166. Importantly the S166A mutant models allows us to dissect the direct effects of PP2A-mediated LSD1 S166 dephosphorylation from transcriptional and epigenetic effects from global PP2A reactivation, targeting range of epigenetic proteins and transcription elongation complex (Aakula et al., 2023; Kauko et al., 2020; Vervoort et al., 2021; Xu et al., 2024).

In summary, identification of S166 phosphorylation as the key regulatory mechanism defining role of LSD1 on epigenetic landscape, resolves a long-standing enigma how this critical epigenetic repressor protein can also mediate gene activation in non-hormonal cancer cells. Our results further demonstrate how tissue microenvironment and immune cell infiltration can be controlled via dephosphorylation of a single phosphorylation site in cancer cell. Collectively our results motivate paying further emphasis on the overlooked power of individual phosphorylation sites in defining not only cell autonomous functions but also controlling tissue homeostasis. Expanding this concept to hundreds of thousands of phosphorylation sites that are yet functionally uncharacterized, it is tempting to speculate that systematic charting of phoshorylation sites at unstructured protein regions may fundamentally change our comprehension of non-genetic control of cancer and other pathological and physiological conditions.

## Material and Methods

### Cell culture

The cell lines used in the study included the KRAS-mutant lung cancer cell lines A549, H358, H441 and H460, HeLa cells and bronchial epithelial cell line HBEC3-KT (HBEC), immortalized with CDK4 and hTERT (Sato et al., 2013). Cells were authenticated using STR profiling by the European Collection of Authenticated Cell Cultures. All cell lines were cultured in a humified incubator with 5% CO2 and a temperature of 37° C. Cells were cultured in media as per the ATCC recommendations. and split once they reached a confluence of around 75%. All cells were consistently tested negative for mycoplasma.

### Transfections

Gene silencing was carried out using the Lipofectamine™ RNAiMAX Transfection Reagent (#13778150), while protein overexpression was achieved with the jetPRIME® transfection reagent Polyplus (#114-15) according to the manufacturer’s instructions.

siRNA sequences and/or provider catalogue #: siCtrl (#1): AllStars Neg. Control siRNA, Cat. No. / ID: 1027281 (Qiagen), siCtrl (#2): CGU ACG CGG AAU ACU UCG A, siHRAS: GAA CCC UCC UGA UGA GAG U, siKRAS: AGA GUG CCU UGA CGA UAC A, siNRAS: GAA AUA CGC CAG UAC CGA A, siSET (#1): UGC AGA CAC UUG UGG AUG G, siSET (#2): AAU GCA GUG CCU CUU CAU C, siLSD1: FlexiTube siRNA #SI02780932 (Qiagen), siSNAI2 (#1): GCU ACC CAA UGG CCU CUC U, siSNAI2 (#2): UCU GGC UGC UGU GUA GCA C, siMYBBP1A (#1): UCU UUC AGU CAG GUC GGC UGG UGA A, siMYBBP1A (#2): GAG GTC CTC AAA GCC GAC TTG AAT A.

### Site directed mutagenesis

LSD1 expressing pcDNA3.1 plasmid was a kind gift from Professor Fan Zhang. The KDM1A S166 mutants were generated through site-directed mutagenesis utilizing polymerase chain reaction (PCR), done with Phusion Green Hot Start High-Fidelity PCR Master Mix (F-5665, Thermo Scientific) and the DpnI enzyme from the QuickChange Lightning Site-Directed Mutagenesis Kit (210518-5, Agilent Technologies). The subsequent LSD1 variants were produced with primers designated as Forward (F) and Reverse (R), documented in the 5′−3′ orientation: S131A (F: C AAC CTC GCG GAA GAT GAG TAT TAT TCA GAA G; R: C ATC TTC CGC GAG GTT GGC CAA GCT TTC AT). S131D: (F: C AAC CTC GAT GAA GAT GAG TAT TAT TCA GAA G; R: C ATC TTC ATC GAG GTT GGC CAA GCT TTC ATC). S137A (F: G TAT TAT GCG GAA GAA GAG AGA AAT GCC AAAG) (R: C TTC TTC CGC ATA ATA CTC ATC TTC TGA GAG G). S137D (F: G TAT TAT GAT GAA GAA GAG AGA AAT GCC AAA G; (R: C TTC TTC ATC ATA ATA CTC ATC TTC TGA GAG G). S166D (F: G AAA ATG AAG ATG AGC CTG AAG AAC CAT CGG; R: C AGG CTC ATC TTC ATT TTC TTC CTC AGG TGG). S166A (F: G AAA ATG AAG CGG AGC CTG AAG AAC CAT CGG) (R: C AGG CTC CGC TTC ATT TTC TTC CTC AGG TGG).

### CRISPR/Cas9-mediated LSD1 mutagenesis

CRISPR-Cas9 genome editing was performed using a non-viral ribonucleoprotein (RNP) delivery system to introduce targeted site-directed mutations in the LSD1 (KDM1A) gene. The approach utilized synthetic crRNA:tracrRNA duplexes and a single-stranded oligodeoxynucleotide (ssODN) donor template to mediate homology-directed repair (HDR). Guide RNA (gRNA) sequences and HDR templates were designed using Benchling (www.benchling.com), ensuring specificity and minimizing off-target effects. Silent mutations were incorporated into the Protospacer Adjacent Motif (PAM) to prevent recutting by Cas9 post-editing. All reagents were ordered from Integrated DNA Technologies (IDT). crRNA: 5’-AAG UGA GCC UGA AGA ACC AUG UUU UAG AGC UAU GCU -3’; HDR Template: GAG AAG GAA AAG AAG CTT CCC CCA CCT CCC CCA CAA GCC CCT CCA GAA GAG GAA AAT GAA GCT GAG CCT GAA GAA CCA TCG GGT GAG TTG TAG TAT CCA ACC ACA GTT CTG TTT TAT CTT AG.

To prepare the CRISPR-Cas9 RNP complex, 120 pmol of crRNA was mixed with an equivalent amount of Alt-R CRISPR-Cas9 tracrRNA (tracrRNA, IDT) in 2.6 μL of CAS9 buffer (20 mM HEPES (pH 7.5), 150 mM KCl, 1 mM MgCl₂, 10% glycerol, and 1 mM TCEP). This mixture was heated at 95 °C for 5 minutes and then allowed to cool at room temperature (RT) for approximately 5–10 minutes. Subsequently, 100 pmol of Alt-R® S.p. HiFi Cas9 Nuclease V3 (IDT) in 3.36 μL of CAS9 buffer was added gradually to the crRNA:tracrRNA duplex. The solution was incubated for 20 minutes at RT to facilitate the formation of the active RNP complex. The assembled RNP complex was combined with 100 pmol of HDR template and introduced into H460 cells (2.5 × 10⁵ per reaction) suspended in Nucleofector SF Cell Line Solution (Lonza, V4XC-2032). A 20 μL aliquot of the final cell/RNP mixture was transferred into a Nucleocuvette Strip (Lonza, V4XC-2032) and transfected using the 4D-Nucleofector™ System (Lonza) under the optimized CM-130 program. Cells were transferred to 6 well plates and grown for 72h. The cells were single-cell cloned and heterozygous knock-in was confirmed by Sanger sequencing.

### Subcellular protein fractionation

Cells were fractionated with the Subcellular Protein Fractionation Kit for Cultured Cells from Thermo Fisher Scientific (#78840) according to the manufacturer’s guidelines. Prior to fractionation, the corresponding buffers were augmented with protease inhibitors included in the kit. Cells were initially harvested using trypsinization, subsequently washed twice in PBS, and 1 million cells were utilized for fractionation. To fractionate the cytoplasmic fractions, cells were suspended in 100 µ l of cytoplasmic extraction buffer (CEB) and treated with gentle agitation for 10 minutes at 4° C. Post-incubation cells were centrifuged at 500g for 5 minutes, and the supernatant containing the cytoplasmic fractions was collected in fresh tubes and the pellets were suspended in100 µ l of membrane extraction buffer (MEB), followed by vortexing at maximum for 5 seconds and subsequent incubation at 4° C for 10 minutes with moderate agitation. The tubes were centrifuged at 3000g for 5 minutes, after which the supernatant containing membrane-bound proteins was transferred to new tubes. The pellets containing the nucleus were initially washed in PBS, thereafter, suspended in 50 µ l of nuclear isolation buffer (NEB), and vortexed for 10 seconds at maximum speed. The tubes were thereafter incubated for 30 minutes at 4° C with agitation. The tubes with lysed nuclei were centrifuged at 5000g for 10 minutes to precipitate the chromatin, while the nucleoplasmic proteins in the supernatant were transferred to new tubes. The chromatin was subsequently treated with 50 µ l of nuclear isolation buffer (NEB) augmented with 150U of micrococcal nuclease (MNAse) and 5mM calcium chloride, followed by vortexing at maximum speed for 10 seconds. The tubes were incubated at room temperature for 15 minutes and vortexed for 15 seconds at maximum speed followed by centrifugation at 16,000g for 10 minutes. The supernatant, comprising chromatin-associated proteins, was transferred to new tubes.

## Western blotting

Cells were lysed using RIPA buffer (50 mM Tris–HCl, pH 7.5, 0.5% DOC, 0.1% SDS, 1% NP-40, and 150 mM NaCl) supplemented with protease and phosphatase inhibitors (#4693159001 and #4906837001; Roche), followed by sonication at maximum intensity with a pulse duration of ± 30 seconds. Following centrifugation at 16,000g for 30 minutes, the lysates were transferred to a new tube, and protein concentration was quantified using the BCA test (Pierce). Sixfold loading buffer was iadded to the lysates, which were subsequently heated at 95°C for 10 minutes. Equal volumes of lysates were applied to 4–20% precast gradient gels (Bio-Rad) and subjected to electrophoresis at 80–120 V. Proteins were transferred onto a PVDF membrane (Bio-Rad) and incubated for 1 hour at room temperature for blocking. Membranes were incubated overnight with the primary antibody and subsequently washed. Detection was performed using HRP-labeled secondary antibodies (DAKO) with subsequent incubation in Pierce ECL Western Blotting Substrate (Thermo Fisher Scientific), followed by detection via ChemiDoc Imaging Systems (Bio-Rad Laboratories). The following antibodies, at the indicated dilutions, were used: FLAG (F3165, 1:1,000; Sigma-Aldrich); GAPDH (5G4-6C5, 1:5,000; HyTest); GFP (sc-9996, 1:500; Santa Cruz Biotechnology); HDAC1 (06-720-25UG, 1:1,000; Sigma-Aldrich) and HDAC2 (sc-9959, 1:1,000; Santa Cruz Biotechnology); H3 (sc-374669 (C-2), 1:1,000; Santa Cruz Biotechnology); LSD1 (ab17721; 1:1000; Abcam), SNAI2 (12129-1-AP, 1:1000, Proteintech), MYBBP1A (14524-1-AP, 1:1000; Proteintech) and SET1 (I2PP2A (F-9), sc-133138, 1:1,000; Santa Cruz Biotechnology)

### Light microscopy setup

The spinning-disk confocal microscope employed was a Marianas spinning-disk imaging system featuring a Yokogawa CSU-W1 scanning unit mounted on an inverted Zeiss Axio Observer Z1 microscope, which was controlled by SlideBook 6 (Intelligent Imaging Innovations, Inc.). Images were captured using a Prime BSI sCMOS camera (chip size 2,048 × 2,048; Photometrics). The objectives utilized were 63x oil (NA 1.4 oil, Zeiss Plan-Apochromat, M27) and 100x oil (NA 1.46 oil, Zeiss Plan-Apochromat, M27).

### Focal adhesion coverage, β1 integrin activation, and stress fiber quantification

Cells were seeded in Ibidi 8-well dishes (Ibidi, 80807), pre-coated with poly-L-lysine (0.01%, 45 minutes at room temperature), followed by bovine plasma fibronectin (10 µ g/mL in PBS, 2 hours at 37°C). After 24 hours of seeding, cells were fixed with 4% formaldehyde in PBS for 10 minutes at room temperature. The cells were then washed three times with PBS and permeabilized using 0.2% Triton X-100 for 5 minutes at room temperature. Following three additional washes with PBS, immunostaining was performed by incubating the cells with primary antibodies, anti-Paxillin (1:400 in PBS) and anti-12G10 (1:15 in PBS), for 1 hour at room temperature in the dark. The cells were subsequently washed three times with PBS and incubated with secondary antibodies: anti-rabbit Alexa Fluor 488 (1:400 in PBS), anti-mouse Alexa Fluor 488 (1:400 in PBS), Alexa Fluor Plus 647 Phalloidin (1:2000), and DAPI (1.5 µg/mL) for 45 minutes at room temperature in the dark. Afterward, the cells were washed three times with PBS and stored at 4°C in Alexa Fluor Plus 647 Phalloidin (1:3000 in PBS) until imaging. Samples were washed three times with PBS just before imaging.

High-resolution images were captured in 3D using a spinning disk confocal microscope. Automatic cell segmentation was performed using the ZeroCostDL4Mic cellpose notebook (Pachitariu and Stringer, 2022; von Chamier et al., 2021) and manually validated in Fiji. For stress fiber quantification, images were processed with a bandpass filter (filter large=20, filter small=4, tolerance=5, autoscale, saturate), and then converted to 8-bit format. Using the Ridge Detection Fiji plugin (doi:10.5281/ZENODO.845874, doi:10.1109/34.659930), stress fibers were identified and quantified with the following parameters: line width=6, high contrast=230, low contrast=20, sigma=2.23, lower threshold=0.17, upper threshold=3.40, and minimum line length=50. The coverage of stress fibers was quantified from the Ridge Detection segmentation output using Fiji’s "Analyze Particles" function in each segmented cell. Focal adhesion number, size, and coverage analyses were performed on carefully chosen z-planes that prominently displayed the focal adhesions. Initially, paxillin-stained images were processed with Fiji’s "Remove Background" feature, applying a 50 px kernel. These images underwent segmentation via a custom-trained random forest classifier using LabKit. The coverage of focal adhesions was quantified from the LabKit segmentation output using Fiji’s "Analyze Particles" function in each segmented cell. Dot plots were generated using PlotsOfData.

The following antibodies, at the indicated dilutions, were used: anti-Paxillin (Abcam, ab32084, diluted for immunofluorescence (IF) at 1:400 in PBS) and 12G10 (a kind gift from Prof. Johanna Ivaska, diluted for IF at 1:15 in PBS). The secondary antibodies employed were Alexa Fluor 488 conjugated (Invitrogen; A11008, A11001), diluted for IF in PBS. The following fluorescent counterstains were used to visualize DNA/nuclei and the actin cytoskeleton: DAPI (4’,6-diamidino-2-phenylindole, dihydrochloride), provided by Thermo Fisher Scientific (D1306), and Alexa Fluor Plus 647 Phalloidin (Invitrogen, A30107). Poly-L-lysine and bovine plasma fibronectin (A-005-M and 341631, respectively) were provided by Merck. Cells were fixed in 4% formaldehyde in PBS (Thermo Fisher Scientific, 28908) and permeabilized in 0.2% Triton-X-100 in PBS (Merck, X100-100ML).

### Proximity Ligation Assay

Duolink PLA kit was used according to the manufacturer’s protocol. The cells were seeded on ibidi 8-well plates. Confluent wells were washed with PBS, then blocked with Duolink Blocking Solution in a humidity chamber at 37°C for 60min. Primary antibodies were diluted 1:300 in Duolink Antibody Diluent and the cells were incubated with the antibodies overnight in 4°C. The wells were washed twice in Wash Buffer A and PLA Probes were diluted 1:5 in the antibody diluent. The wells were incubated with the probes in a humidity chamber at 37°C for 1h. The wells were washed twice in Wash Buffer A. Ligase was diluted 1:40 in Ligation Buffer and the wells were incubated for 30 minutes at 37°C. The wells were washed twice and incubated for 100 minutes at 37°C with Polymerase diluted 1:80 in Amplification Buffer. The wells were washed twice in Wash Buffer B and mounted with Duolink in Situ Mounting Medium with DAPI. The cells were imaged with Zeiss LSM880 confocal microscope. The abtibodies used were as follows: SET (I2PP2A (F-9), sc-133138) and FLAG (F3165, 1:1,000; Sigma-Aldrich).

### Incucyte growth assay

H460 cells WT clone #1 and S166A clone #1 were seeded into a 96-well plate (3x103 cells per well). The next day, the cells were treated with DMSO or Panobinostat (50 nM) (12 wells/condition). The cells were treated with drugs twice per week followed two-week drug holiday. The confluency of the wells was determined daily using an IncuCyte ZOOM live-cell analysis system (Essen Bioscience, Royston Hertfordshire, UK).

### Histone PTMs mass spectrometry analysis

Histones were enriched from 5x106 H460 cells as previously described (Noberini et al., 2020). Approximately 4 µg of histone octamer were mixed with an equal amount of heavy-isotope labelled histones, which were used as an internal standard (super-SILAC mix)(Noberini et al., 2023), and separated on a 17% SDS-PAGE gel. Histone bands were excised, chemically acylated with propionic anhydride and in-gel digested with trypsin, followed by peptide N-terminal derivatization with phenyl isocyanate (PIC) (Noberini et al., 2021). Peptide mixtures were separated by reversed-phase chromatography on an EASY-Spray column (Thermo Fisher Scientific), 25-cm long (inner diameter 75 µm, PepMap C18, 2 µm particles), which was connected online to a Q Exactive HF instrument (Thermo Fisher Scientific) through an EASY-Spray™ Ion Source (Thermo Fisher Scientific), as described (Noberini et al., 2021). The acquired RAW data were analyzed using EpiProfile 2.0 (Yuan et al., 2018), selecting the SILAC option, followed by manual validation. For each histone modified peptide, the % relative abundance (%RA) for the sample (light channel - L) or the internal standard (heavy channel - H) was estimated and Light/Heavy (L/H) ratios of %RA were then calculated.

### Quantitative RT-PCR

RNA was extracted utilizing the NucleoSpin RNA kit (Macherey-Nagel) and subsequently reverse-transcribed into cDNA employing random primers, dNTP mix (Thermo Fisher Scientific), and the Promega cDNA kit (M3681) M-MLV Reverse Transcriptase, RNase H Minus, Point Mutant, in accordance with the manufacturer’s instructions. To amplify the target genes, AccuTarget ™ qPCR Screening kit (Bioneer) containing a pre-designed primer set for each target gene in a 96 well plate format was used. The PCR amplification carried out with the PowerUp SYBR Green Master Mix (Thermo Fisher Scientific) in the QuantStudio 12K Flex Real-Time PCR System (Thermo Fisher Scientific). Gene expression levels were normalized to the housekeeping gene GAPDH, and the 2−ΔΔCT method was employed to calculate these values.

### Protein expression and purification

Full-length LSD1 (in pET28a vector) and CoREST (residues 308-485) were co-expressed in E. coli (BL21(DE3) strain) using a pCDF DUET1 plasmid as described earlier (Astro et al., 2022). Both proteins had a His-SUMO tag at their N-terminus. Cells were grown in LB medium at 37 °C and 200 rpm until OD600 = 0.8. Protein expression was induced overnight by the addition of 0.25 mM isopropyl β-D-1-thiogalactopyranoside at 17 °C. Cells were harvested at 5000 rpm, 10 °C for 10 min, flash-frozen in liquid nitrogen, and stored at -80 °C. Cells were disrupted by sonication, and the suspension was clarified by centrifugation at 56,000 × g, 4 °C for 50 min. Pellets were re-suspended in 50 mM NaH2PO4 pH 8.0, 300 mM NaCl, 5% glycerol, and 7.5 mM imidazole. They were then loaded onto a HiTrap nickel affinity chromatography. LSD1-CoREST eluted at 500 mM imidazole and was dialyzed overnight with the addition of His-tagged SUMO protease. The protein complex was finally gel-filtered through a Superdex 200 10/300 column equilibrated in 25 mM NaH2PO4 pH 7.2, 5% glycerol. The concentration of LSD1/CoREST is estimated by measuring the flavin absorbance at 458 nm (ε = 10790 M⁻ ¹cm⁻ ¹).

Recombinant expression of MYBBP1A was carried out by Turku Protein Core (TuProtCore, University of Turku and Åbo Akademi University) in E. coli Rosetta(2)DE3 pLysS cells (Novagen), following the manufacturers guidelines. Protein purification was carried out through a polyHis-tag affinity chromatography with a HisTrap HP column (Cytiva), followed by a TEV protease (New England BioLabs) cleavage to remove the tag. A final clean-up was carried out with a size exclusion Superdex 200 Increase 10/300 GL, equilibrated with the final storage buffer (50 mM NaH2PO4, 300 mM NaCl, pH 7.4).

### LSD1 activity assays

Peptides were purchased from Genscript. All other reagents were from Merck. Activity measurements were performed with a peroxidase-coupled assay using a Clariostar plate reader (BMG Labtech). The reactions were carried out in 50 mM HEPES pH 8.5, 0.1 mM amplex red, and 0.3 mM horseradish peroxidase using 0.3 µM LSD1-CoREST. The substrate peptide comprising residue 1-21 of H3 dimethylated at Lys4, was serially diluted from 40 µM to 0.31 µM. The measured fluorescence signal reflects the enzymatic conversion of amplex red to resorufin. Non-linear regression analysis using GraphPad Prism software was used to determine the kcat and KM values.

### Differential Scanning Fluorescence (nanoDSF) analysis

The thermal stability of LSD1 WT and S166A mutant was assessed using differential scanning fluorescence (nanoDSF) on the Prometheus NT.48 system (NanoTemper Technologies). This technique monitors shifts in the fluorescence emission of tryptophan (Trp) residues as a function of temperature, allowing for the determination of the melting temperature (Tm) and protein aggregation onset. Protein samples were prepared at a concentration of approximately 0.2 mg/mL in a uniform buffer condition. For each variant, 10 µL of protein solution was loaded into High Sensitivity nanoDSF capillaries (NanoTemper Technologies). The samples were heated from 20°C to 95°C at the rate of 1°C/min and fluorescence emission was measured at 350 nm and 330 nm (F350/F330 ratio). The F350/F330 ratio was plotted against temperature to determine the inflection point (IP), which corresponds to the melting temperature (Tm). Aggregation onset was determined based on light scattering measurements.

### Targeted proteomics assay

The assay was performed by the Proteomics core facility of the Turku Centre for Biotechnology. Cells were lysed and denatured in 8 M urea in 50 mM Tris-HCl, pH 8 with protease and phosphatase inhibitors. Samples were reduced with 10 mM D,L-dithiothreitol and alkylated with 40 mM iodoacetamide. Samples were digested overnight with sequencing grade modified trypsin (Promega). After digestion peptide samples were desalted with a Sep-Pak tC18 96-well plate (Waters) divided into total protein and phosphopeptide samples and evaporated to dryness. Phosphopeptides were enriched with Thermo Scientific High Select Fe-NTA phosphopeptide enrichment kit according to manufacturer instructions and evaporated to dryness. Samples were dissolved in 0.1% formic acid and for each selected peptide, 1 pmol of isotopically labelled peptide (13C6,15N4-Arginine) was spiked in the samples and used as an internal standard for quantification by Parallel reaction monitoring (PRM). Samplee were analyzed by PRM using an Orbitrap Exploris 480 mass spectrometer (Thermo Fisher Scientific) coupled to an EASY-nanoLC 1200 UPLC system (Thermo Fisher Scientific). Peptides were first loaded on a trapping column and subsequently separated inline on a 15 cm C18 column (75 μm x 15 cm, ReproSil-Pur 3 μm 120 Å C18-AQ, Dr.

Maisch HPLC GmbH, Ammerbuch-Entringen, Germany). The mobile phase consisted of water with 0.1% formic acid (solvent A) and acetonitrile/water (80:20 (v/v)) with 0.1% formic acid (solvent B). A 60 min gradient was used to elute peptides (50 min from 5 % to 36 min solvent B and in 5 min from 39% to 100% of solvent B, followed by 5 min wash stage with solvent B). The Orbitrap Exploris 480 was operated in positive ionization mode with an EASY-Spray nanosource at 2 kV and at a source temperature of 280 °C. A PRM method was used for data acquisition to target all selected peptides with a quadrupole isolation window set to 1.6 m/z with detection in the Orbitrap mass analyser at a 30 K resolution. HCD fragmentation was used for MS2 fragmentation with normalized collision energy of 30%, normalized AGC was set at 300% and the maximum injection time was 52 ms. All data was acquired with XCalibur software v.4.6. Acquired data were analyzed with the Skyline software (v24.1.1.398) to identify peptides and quantify peak areas. Peak areas were normalized to internal standard peptides and the ratio of peak areas was used to compare the relative abundance of peptides. The human protein database was used as the background proteome.

### Proteomic analysis of Xenograft samples

Human xenograft samples from mice were powdered using Covaris CP01 Manual Pulverizer according to manufacturer instructions. Powdered samples were lysed in 5 % SDS in 100 mM triethylammonium bicarbonate buffer (TEAB) and sonicated with Bioruptor Plus (Diagenode) sonicator using 15 x 30 s on/off cycles. Samples were reduced with 10 mM Tris (2-carboxyethyl) phosphine and alkylated with 40 mM chloroacetamide. Samples proteins were aggregated to MagResyn Hydroxyl beads (ReSyn Bioscience) using protein aggregation capture method (Batth et al., 2019) and digested overnight with sequencing grade modified trypsin (Promega), and Lys-C (Wako). After digestion peptide samples were acidified and divided into two: a total protein sample and sample for phosphopeptide enrichment. Total protein samples were acidified to 1 % trifluoroacetic acid concentration (pH 2) and desalted with a Sep-Pak tC18 96-well plate (Waters) and evaporated to dryness. Samples for phosphopeptide enrichment were evaporated to dryness and phosphopeptides were enriched with High Select Fe-NTA Phosphopeptide Enrichment Kit (Thermo Fisher Scientific) according to manufacturer instructions. For total protein DIA analysis 800 ng of peptides were injected and analyzed. For phosphopeptide DIA analysis half of the samples were injected and analyzed.

The LC-ESI-MS/MS analysis was performed on a nanoflow HPLC system (Easy-nLC1200, Thermo Fisher Scientific) coupled to the Orbitrap Lumos Fusion mass spectrometer (Thermo Fisher Scientific, Bremen, Germany) equipped with a nano-electrospray ionization source and FAIMS interface. Compensation voltages of -50 V and -70 V were used. Peptides were first loaded on a trapping column and subsequently separated inline on a 15 cm C18 column (75 μm x 15 cm, ReproSil-Pur 3 μm 120 Å C18-AQ, Dr. Maisch HPLC GmbH, Ammerbuch-Entringen, Germany). The mobile phase consisted of water with 0.1% formic acid (solvent A) or acetonitrile/water (80:20 (v/v)) with 0.1% formic acid (solvent B). A 120 min gradient was used to elute peptides (62 min from 5 % to 21 % solvent B followed by 48 min from 21 % to 36 % solvent B and in 5 min from 36% to 100% of solvent B, followed by 5 min wash stage with solvent B).

Samples were analyzed by a data independent acquisition (DIA) LC-MS/MS method. MS data was acquired automatically by using Thermo Xcalibur 4.6 software (Thermo Fisher Scientific). In a DIA method a duty cycle contained one full scan (400 -1000 m/z) and 30 DIA MS/MS scans covering the mass range 400 -1000 with variable width isolation windows.

DIA data analysis consisted of protein identifications and label free quantifications of protein abundances. Data was analyzed by Spectronaut software (Biognosys; version 18.7.240325). DirectDIA approach was used to identify proteins and label-free quantifications were performed with MaxLFQ. The PTM analysis was performed on phosphopeptide enriched samples. Main data analysis parameters in Spectronaut were: Trypsin was selected as enzyme and two missed cleavages were allowed. Carbamidometholation was set as fixed modification and N-terminal acetylation and methionine oxidation were set as variable modifications. Search was performed against Swiss-Prot Homo Sapiens database (Uniprot 2024-01 release) and Universal Contaminant database (Frankenfield et al., 2022). FDR value of 0.01 was used on PSM, Peptide and Protein Group levels. Quantification was done on MS2 level and area was selected as quantification type. Normalization was performed using local normalization setting. For PTM analysis PTM localization probalibily cutoff was set as 0.75.

### Affinity purification mass spectrometry (AP-MS)

H460 cells were seeded in 10 cm plates at a density of 1 million cells per plate. Following day cells were transfected with either pcDNA3.1 empty vector, or wild type or 166A mutant LSD1plasmids using the jetPRIME® transfection reagent Polyplus (#114-15). 48h prior to transfection immunoprecipitation was done using the Pierce™ MS-Compatible Magnetic IP Kits (90409). Briefly, cells were collected by scraping in 500 µ lof lysis buffer and incubation at 4°C for 20 minutes. Lysates were collected by centrifugation at full speed for 20 minutes and the protein concentration was determined using BCA assay. From each sample, 10 µg of protein was collected as input while 1000 µg was incubated overnight at 4°C, with 5 µg Monoclonal ANTI-FLAG® M2 antibody (F3165-1MG, Sigma). The antigen-antibody mix from the overnight incubation was added to 20 µl of prewashed Protein A/G Magnetic Beads and rotated at room temperature for 60 min. Beads were washed and the protein complexes were eluted by incubating the beads in 100μL of IP-MS Elution Buffer. The elutant was dried in a speed vacuum concentrator and the samples were further processed for MS analysis.

Samples were dissolved in 0.1% formic acid and analyzed using a Q Exactive HF mass spectrometer (Thermo Fisher Scientific) coupled to an EASY-nLC 1200 UPLC system (Thermo Fisher Scientific). Peptides were first loaded on a trapping column and subsequently separated inline on a 15 cm C18 column (75 μm x 15 cm, ReproSil-Pur 3 μm 120 Å C18-AQ, Dr. Maisch HPLC GmbH, Ammerbuch-Entringen, Germany). The mobile phase consisted of water with 0.1% formic acid (solvent A) and acetonitrile/water (80:20 (v/v)) with 0.1% formic acid (solvent B).

A 60 min gradient was used to elute peptides (50 min from 6 % to 39 min solvent B and in 5 min from 39% to 100% of solvent B, followed by 5 min wash stage with solvent B).

MS data was acquired automatically by using Thermo Xcalibur 4.1 software (Thermo Fisher Scientific). A data dependent acquisition method consisted of an Orbitrap MS survey scan of mass range 300–1750 m/z followed by HCD fragmentation for the most intense peptide ions.

The mass spectrometer was operated in the positive ionization mode, with the nano-spray voltage set at 2.1 kV and the source temperature at 300 °C. The acquisition was performed in the data-dependent acquisition (DDA) mode and full mass spectrometry (MS) scans, with 1 microscan at a resolution of 120,000, were used over an m/z range of 350–1750, with detection in the Orbitrap mass analyzer. The auto gain control (AGC) was set to 3e6 and the injection time to 100 ms. In DDA analysis from each MS1 survey scan, ten most intense ions above a threshold were selected for fragmentation. Fragment ion spectra were produced via high-energy collision dissociation (HCD) at normalized collision energy of 27 and acquired in the Orbitrap mass analyzer at resolution 15,000. The AGC and injection time were set to 50,000 ms and 200 ms, respectively, and an isolation window of 1.8 m/z was used.

Data files were searched for protein identification using Proteome Discoverer 3.1 software (Thermo Fisher Scientific) connected to an in-house server running Mascot 2.8.3 software (Matrix Science). The search was conducted against a database for Homo sapiens and common laboratory contaminants. For peptide identification, a precursor ion mass tolerance of 10 ppm was used for MS1 level, trypsin was chosen as enzyme and up to two missed cleavages were allowed. The fragment ion mass tolerance was set to 0.02 Da for MS2 spectra. Oxidation of methionine and N-terminal protein acetylation were set as variable modifications, and carbamidomethylation on cysteine residues was set as a fixed modification. The false discovery rate (FDR) in peptide identification was set to a maximum of 5% for medium confidence peptides and maximum of 1% for high confidence peptides.

### Co-immunoprecipitation analysis

Cells were seeded in 10 cm plates at a density of 1 million cells per plate and cultured according to growth conditions. Following day, the cells were transfected with the corresponding plasmids utilizing the jetPRIME® transfection reagent Polyplus (#114-15). After 48h cells were harvested by scraping on ice and subsequently lysed with a solution containing 100 mM NaCl, 1 mM MgCl2, 10% glycerol, 0.2% protease inhibitor tablet (Roche), and 25 units/ml Benzonase (Millipore). The tubes containing cell lysates were incubated for 15 minutes at 4° C with rotation, after which the concentrations of NaCl and EDTA were raised to 200 mM and 2 mM, respectively and an additional 10-minute incubation under similar conditions was continued. The lysates were subsequently centrifuged at maximum speed, and the supernatant was collected in fresh tubes. Protein concentration was determined using BCA assay and 10% of the supernatant was collected in new tubes as input. The residual supernatant was utilized for immunoprecipitation. Lysates were introduced into the tubes containing prewashed ChromoTek GFP-Trap® Agarose (#gta-20) or ChromoTek DYKDDDDK Fab-Trap™ Agarose (#ffa) beads. Protein interactions were enhanced by rotating the tubes for 90 minutes at 4° C. The beads were rinsed three times in lysis buffer and subsequently boiled at 95° C in 2X lysis buffer. Protein interactions were assessed using western blotting.

### In-vitro pull down assay

To determine direct protein interaction in vitro, 1 µ g of purified proteins were incubated together for one hour in total volume of 150 ul (37 °C; 300rpm). Prior to incubation 10% of the input was collected in new tubes and the remaining lysates was added to 10 µ l of prewashed beads. The total volume of the reaction was made to 500 µ l using buffer and the tubes were rotated for 60 minutes at room temperature. Beads were washed 4 times in 500 µ l of buffer and subsequently boiled at 95° C in 2X lysis buffer. Protein interactions were assessed using western blotting.

### RNA sequencing

RNA sequencing was performed at the Finnish Functional Genomic Center, Turku Bioscience Centre, University of Turku and Åbo Akademi Univeristy. RNA was extracted utilizing the NucleoSpin TriPrep kit (Macherey-Nagel), subsequently treated with DNase to eliminate genomic DNA. The quality of the samples was ensured using Agilent Bioanalyzer 2100 or Advanced Analytical Fragment Analyzer. Sample concentration was measured with Qubit®/Quant-IT® Fluorometric Quantitation, Life Technologies, and/or Nanodrop ND-2000, Thermo Scientific. Library preparation was performed according to the Illumina Stranded mRNA Preparation, Illumina (1000000124518). Sequencing run was performed using Illumina NovaSeq 6000 S1 v1.5 instrument with a read length of 2 x 50 bp. Raw sequencing reads were assessed for quality using FastQC (v0.11.8) and summarized using MultiQC (v1.10). The quality of the reads was evaluated based on standard metrics, including per-base sequence quality, sequence length distribution, and GC content. Only high-quality reads were retained for downstream analysis. Filtered reads were aligned to the human reference genome (hg38) using Rsubread (v2.0.0) within the Bioconductor (v3.9) framework in R (v3.6.1). The RefSeq gene annotation provided in Rsubread was used for feature mapping. The alignment was performed in a paired-end mode with annotation-based mapping enabled. Gene-level quantification of aligned reads was performed using the featureCounts function from Rsubread. The assignment was done based on the in-built RefSeq gene annotation, considering strand-specificity (reverse-stranded mode). Only reads that mapped uniquely to known genes were counted. Normalization of read counts was performed using the edgeR (v3.28.0) package to calculate counts per million (CPM) and reads per kilobase per million mapped reads (RPKM). Differential expression analysis was conducted using ROTS (v1.14.0), comparing the experimental group (166A) to the control group. Genes with a false discovery rate (FDR) below 0.05 and a fold-change ≥2 were considered significantly differentially expressed. To evaluate sample relationships, hierarchical clustering was performed using amap (v0.8-18) with Pearson correlation as the distance metric. Principal component analysis (PCA) was conducted using the pca function in R, with unit variance scaling applied. Correlation analysis between samples was computed using the Pearson method. Differentially expressed genes (DEGs) were visualized using multiple graphical representations, including MA plots, volcano plots, heatmaps, PCA plots, and swarm plots. These visualizations were generated to aid in interpretation and quality control of the differential expression analysis.

### Assay for Transposase-Accessible Chromatin using sequencing (ATAC-seq)

ATAC-seq was done, according to the published protocol (Buenrostro et al., 2015). Library sizes were ascertained using fragment analysis, followed by 2 × 50 paired-end sequencing conducted on the Illumina NovaSeq 6000 SP v1.5. Quality assurance of ATAC-seq sequencing libraries were conducted with FastQC version 0.11.9. Bowtie2 version 2.4.5 were used to align the reads on the human genome GRCh38. Reads were converted to bam with samtools version 1.8, sorted with Picard SortSam version 2.27.1 and indexed with Picard BuildBamIndex version 2.27.1. Duplicated reads were marked with Picard MarkDuplicates version 2.27.1.

Peaks were called with MACS2 v2.2.9.1 with the following parameters: --gsize hs –qvalue 0.05 –format bam –nomodel –bdg –call-summits. Homer version 4.10 was used to annotate the peak summits by running the annotatePeak.pl against hg38 annotation. Standardized peaks were computed starting from peak summit using the bedtools version 2.29.1 slop command with parameters: -g $GENOME -l 250 -r 249. Bedtools version 2.29.1 coverage command was used to compute the read counts of standardized peaks. The standardized peak read count was normalized using the median of ratio normalization and then converted to RPM. Log2 fold change was computed from mean RPM values comparing signal intensities of corresponding genomic locations between LSD1 S166A mutant and WT samples. Differentially accessible regions were determined using DESeq2 version 1.40.2. Heatmaps were generated using R version 4.3.1 and R-package pheatmap 1.0.12 and volcano plots were generated using R-package EnhancedVolcano 1.18.0. Footprint analysis was performed using TOBIAS (v0.15.1), specifically using the ATACorrect, ScoreBigwig, and BINDetect functions (Bentsen et al., 2020).

### Reduced Representative Bisulphite Sequencing (RRBS)

DNA was isolated using NucleoSpin TriPrep kit (Macherey-Nagel), and the quality of the samples was ensured using Agilent Bioanalyzer 2100 or Advanced Analytical Fragment Analyzer. Sample concentration was measured with Qubit®/Quant-IT® Fluorometric Quantitation, Life Technologies, and/or Nanodrop ND-2000, Thermo Scientific. Library preparation was performed according to the library preparation protocol, Reduced representation bisulfite sequencing(Boyle et al., 2012). Sequencing run was performed using Illumina NovaSeq 6000 SP v1.5 with a read length of 2 x 50 bp. Quality of RRBS-seq data was assessed with FastQC version 0.11.9. Reads were trimmed with TrimGalore! version 0.6.7 running on top of Cutadapt version 4.2 with the following parameters: --quality 22 --phred33 --gzip --rrbs --fastqc --paired. Trimmed reads were aligned with Bismark version 0.22.3 using the Bowtie2 version 2.5.2 and the following parameters: -- unmapped --ambiguous --ambig_bam --nucleotide_coverage --fastq. Data were aligned to the Homo sapiens GRCh38 genome and Gencode v41 annotations were used. Methylation calls were extracted with the bismark_methylation_extractor and the following parameters: --paired-end -- comprehensive --gzip --bedGraph --remove_spaces --buffer_size 80% -- cytosine_report --ignore_r2 2. Incomplete conversions were filtered out with the filter_non_conversion tool from Bismark. Differential methylation analysis was run using Bioconductor R-package methylKit (Akalin et al., 2012) version 1.26.0 running on R version 4.3.1. For the methylKit analysis, two scenarios were considered: CpG context and tiled window context. For differential methylation calls, the following parameters were set: methylation difference 25% and adjusted q-value 0.01. For the tiled analysis also window size and step size were set to 500 bp to generate tiling, non-overlapping windows. Bioconductor R-package biomaRt version 2.56.1 was used for annotation. R-package pheatmap version 1.0.12 was used for visualization of the methylation levels. Volcano plots were generated using R-package EnhancedVolcano 1.18.0.

### In vivo studies

All animal experiments were approved by the National Animal Experiment Board of Finland (ESAVI/11684/2021 license), and the studies were performed according to the instructions provided by the Institutional Animal Care and Use Committees, University of Turku, Finland.

4–6 weeks of age female athymic nude mice (Hsd:Athymic Nude-Foxn1nu) were purchased from Envigo, Gannat, France. The mice were housed in individually ventilated cages (IVC, Techniplast; up to five mouse/cage), under controlled conditions of light (12-h light/12-h dark), temperature (21±3°C), and humidity (55±15%) in specific pathogen-free conditions at the Central Animal Laboratory, University of Turku (Turku, Finland). The mice were given an irradiated soy protein-free diet (2920X; Envigo—Teklad Diets) and autoclaved tap water ad libitum.

Subcutaneous (s.c.) xenografts were generated by engrafting 1,5x106 H460 cells WT clone #1 and S166A clone #1 suspended in a 150 µL solution of PBS/Matrigel (1: 1 v/v). Tumor growth of H460 s.c. xenografts was monitored two times a week using caliper measurements, and the animals were weighed once a week. The volume of the s.c. tumors was calculated according to the following formula: W2 - L/2 (W—shorter diameter, L—longer diameter of the tumor). Mice were sacrificed when the tumors reached their maximum size (longer diameter reached 15 mm). The collected tumors were divided into two parts: one part was snap-frozen in liquid nitrogen, and the other was fixed in 10% formalin for two days. The formalin-fixed paraffin-embedded (FFPE) blocks were prepared at the Histology core facility, Institute of Biomedicine, University of Turku, Finland.

### Multiplex immunofluorescence (mIF) staining

FFPE blocks were cut to 5µm slides at the Histology core facility, Institute of Biomedicine, University of Turku, Finland. The following antibodies and dyes were used: F4/80 (D2S9R) XP (Cat#70076, CellSignaling), Vimentin (Cat#10366-1-AP, Proteintech), Ly6G (Cat#87048T, CellSignaling), CF555 yellow (92171, Biotium), CF488 green (92171, Biotium), CF640R far-red (92175, Biotium), ProLong Gold Antifade Mountant with DAPI (P36935, P36931, Thermo Scientific).

The multiplex immunofluorescence staining was done using the Leica BOND RX Fully Automated Research Stainer (Leica Biosystems, Germany). Slides were deparaffinised by baking at +60°C for 30 min, followed by incubation in BOND Dewax Solution (AR9222, Leica Biosystems) and absolute ethanol. Slides underwent heat-induced epitope retrieval (HIER) in BOND Epitope Retrieval Solution 2 (pH 9.0; AR9640, Leica Biosystems) for 20 min at +100°C and endogenous peroxidase was blocked by using 3% hydrogen peroxide for 10 min. Prior to primary antibody incubation (20 min, Table X), slides were blocked with Normal Antibody Diluent for 10 min (BD09-999-IL, Immunologic, The Netherlands), followed by application of a polymeric HRP-conjugated secondary antibody (DPVR110HRP-IL, Immunologic) for 10 min. Slides were incubated for 10 min with appropriate fluorophore-conjugated tyramide (CF-dye tyramides, Biotium, CA, USA) in 2 nM final concentration in Tyramide Amplification Buffer Plus (Biotium), supplemented with fresh 0.0015% hydrogen peroxide immediately prior to use. Slides were subjected to HIER in BOND Epitope Retrieval Solution 2 (10 min at +98°C) to strip the primary and secondary antibodies bound to the tissue and the steps from primary antibody incubation were repeated until all markers were labelled. Slides were washed with BOND RX Wash Solution in between steps (Bond Wash Solution, Leica Biosystems) and mounted in ProLong Gold Antifade Mountant with DAPI (P36935, Thermo Scientific, MA, USA). Slides were scanned using a Pannoramic 250 Flash scanner (3DHISTECH, Budapest, Hungary), and images were acquired using a SLIDEVIEWER v2.6 (3DHISTECH). The representative images were collected using Nikon Eclipse Ti2-E widefield inverted microscope using Nikon NIS Elements 4.11 acquisition software.

### Chromatin immuno-precipitation-sequencing (ChIP-seq)

Cells were cultured under standard conditions, and about 2 x 10^8^ cells from each condition were crosslinked using 1% formaldehyde (37%) for 10 minutes at room temperature on a shaker. The reaction was quenched by adding glycine to a final concentration of 0.125M, followed by 5 minutes of incubation at room temperature.

Cells were then washed twice with ice-cold PBS, resuspended, and centrifuged at 4°C to remove supernatant before proceeding to lysis. Lysis was performed using SDS buffer (50 mM Tris-HCl pH 8.0, 0.5% SDS, 100 mM NaCl, 5 mM EDTA, 0.02% NaN3), supplemented with protease inhibitors (complete mini cocktail, Roche). Lysates were incubated at room temperature for 2 minutes before being either stored at -80°C overnight. Chromatin was diluted in half the volume of Triton dilution buffer (100 mM NaCl, 100 mM Tris-HCl pH 8.5, 5 mM EDTA, 5% Triton X-100, 0.02% NaN3) and transferred to Bioruptor® Pico microtubes (Diagenode). Sonication was performed using the Diagenode Bioruptor® Pico at 4°C, applying 45 cycles (30 seconds each) to obtain fragment sizes between 100-300 bp.

Fragmentation was confirmed by agarose gel electrophoresis following DNA purification. Pre-clearing was carried out using BSA-blocked Protein A Sepharose beads, which were washed twice in IP buffer and incubated with chromatin on a rotating wheel at 4°C for 2 hours. Chromatin was quantified using Bradford protein assay (595 nm absorbance, Beckman Coulter Reader) and 2.5% of the input sample was collected and stored at 4°C. For each IP reaction, 1500 μg of chromatin was incubated overnight with 10 μg LSD1 antibody (Abcam, no. 17721) at 4°C on a rotating wheel. As a negative control, an equal amount of IgG (Rabbit) was used for mock samples. Pre-blocked Protein G/A magnetic beads (30 μL per IP) were then added and incubated for 2 hours at 4°C to capture immune complexes. Samples were washed sequentially with four different buffers supplemented with 1x Protease Inhibitor Cocktail (Roche, 04693116001): three washes in mixed micelle buffer (Sucrose 5.2% (w/v), 1% Triton X-100, 20 mM Tris-HCl (pH 8.1), 150 mM NaCl, 0.2% SDS, 5 mM EDTA pH 8.0, 0.02% NaN3,), three washes in buffer 500 (500 mM NaCl, 0.01% Triton X-100, 50 mM HEPES pH 7.9, 0.1% Sodium Deoxycholate, 1 mM EDTA pH 8.0, 0.02% NaN3), followed by three washes in LiCl detergent solution (250 mM LiCl, 0.5% Sodium Deoxycholate,0.5% NP-40, 1 mM EDTA pH 8.0, 10 mM Tris-HCl (pH 8.0), 0.02% NaN3), and finally twice in 1X TE buffer. After washing, chromatin was eluted in 200 μL decrosslinking buffer (0.1 M NaHCO3 and 1% SDS) and incubated overnight at 65°C with 0.25 mg/ml Proteinase K in TE 1X). DNA was purified using the Qiagen PCR purification kit and quantified using the Qubit DNA HS assay (Thermo Fisher Scientific). DNA integrity and yield were assessed before proceeding to qPCR validation and/or ChIP-seq library preparation. The primer sequences used for qPCR ChIP validation designated as Forward (F) and Reverse (R), documented in the 5′−3′ orientation are as follows: GAPDH (F: TTC GCT CTC TGC TCC TCC TG; R: CCT AGC CTC CCG GGT TTC TC), Gene Desert (F: AGC TAT CTG TCG AGC AGC CAA G; R: CAT TCC CCT CTG TTA GTG GAA GG), JUN (F: TCT TCT CTT GCG TGG CTC TC; R: GAG ACA AGT GGC AGA GTC CC), HOXD1 (F: GTG CCA GGC ATT GAG AAA AT; R: GGA ACA GAA TTT GTA CCA GGA GA), NDUFA3 (F: GGG GCC TCG CTG TAA TTC TG; R: GAC GGG CAC TGG GTA GTT G), OSCAR (F: AAC CCG CTT GGA GAT TTG G; R: GAG GAC ACA TCC CGG AAG AG).

### ChIP-seq data analysis

ChIP-seq data were analyzed using the nf-core/chipseq pipeline (https://nf-co.re/chipseq/2.1.0/), aligning to the hg38 genome, with narrow peak calling parameter and a MACS2 p-value threshold of 0.001. The peaks obtained with MACS2 were further filtered by intersecting the peaks identified as common between the two different clones of both the mutant and the control, with the peaks identified as differentially enriched in the mutant or control using the test_differential_abundance function of tidybulk (p-value < 0.05 and |log2FC| > 0.5) on the consensus peak set. Density plots and heatmaps were generated using the deepTools package (v3.4.3). Gene ontology analysis was performed using Enrichr. Regions-to-genes association was performed with GREAT (https://great.stanford.edu/).

### Bioinformatics Analysis

Functional analysis, including Molecular Signature Analysis, Gene Ontology (GO) Term Enrichment Analysis, and Upstream Gene Analysis, was conducted using iPathwayGuide™ (AdvaitaBio Corporation, https://ipathwayguide.advaitabio.com/) or Enricher pathway analysis.

Transcription Factor Enrichment Analysis was performed using the ChEA3 Transcription Factor (TF) Analysis: a gene-set-based transcription factor enrichment tool. To compare the transcriptional landscape of LSD1 S166A mutant and WT clones, a gene set perturbation analysis was performed. The rank plot was generated using GSEA-based similarity scoring (Guo et al., 2023). Integrative multi-omics approach was applied, combining transcriptional profiling (RNA-seq), chromatin accessibility analysis (ATAC-seq), and DNA methylation analysis (RRBS) using Metascape (Zhou et al., 2019).

Protein Structure and Interaction Analysis was performed using the tools as follows: structural modeling of proteins was performed using AlphaFold to predict 3D structures and conformational dynamics]; Sequence-based functional annotation and network-based prediction of protein interactions were performed using SCANNet database. Protein-protein interaction (PPI) networks were analyzed using STRING database (Search Tool for the Retrieval of Interacting Genes/Proteins).

To evaluate the evolutionary conservation of LSD1 phosphorylation sites, the phosphoELM database was utilized. The phosphorylation dynamics of LSD1 at serine 166 (S166) across various human tissues and cell lines were analyzed using the qPTM database (Yu et al., 2023). Intrinsic disorder propensity and surface accessibility of LSD1 were predicted to assess the structural flexibility and solvent exposure of key phosphorylation sites using the eukaryotic phosphorylation site database (EPSD) (Lin et al., 2021). Proteomics datasets to analyse the S166 phosphorylation status in lung adenocarcinoma was integrated from the Cancer Proteome Atlas (CPTAC) by ULCAN (Chandrashekar et al., 2022) using the Apollo dataset (Soltis et al., 2022).

### Statistical analyses

The statistical test used to evaluate the data is described in the respective method section or figure legend and were conducted by MS, OD, EC, RH or GJ . All experiments were conducted a minimum of three times, or as specified in the figure legends. GraphPad Prism was used to perform the statistical analyses except for immunofluoresence analysis of integrin activation, focal adhesions and stress fibers for which statistical analyses randomization tests and Cohen’s D values were computed using code available at (https://github.com/CellMigrationLab/Plot-Stats).

## Acknowledgements

This study was supported by the Jane and Aatos Erkko Foundation (to J.W), Sigrid Juselius Foundation (to J.W, G.J), the Cancer Society of Finland (Syöpäjärjestöt; to J.W, G.J), and the Solutions for Health strategic funding to Åbo Akademi University (to G.J.), the InFLAMES and GeneCellNano Flagships Programme of the Research Council of Finland /RCF) and other RCF projects (decision numbers: 337530, 337531, 357910, 35791, 337120, 352818, 338537), and the Aamu Foundation (A.A and O.D). Professor Johanna Ivaska is greatly acknowledged for her valuable comments and for providing the 12G10 antibody. Professors Fan Zhang and Kenneth Zaret are warmly thanked for providing the LSD1 and MYBBP1 expression vectors, respectively. Professor Shay is thanked for providing the immortalized HBEC cell line. We are very grateful for the services, instrumentation and expert help from following Turku Bioscience Centre (University of Turku and Åbo Akademi University) core facilities all supported by Biocenter Finland: The Cell Imaging and Cytometry Core facility, Turku Proteomics Facility, Finnish Functional Genomics Centre Medical Bioinformatics Centre, and Screening unit. Further the facilities and expertise of the Structural Bioinformatics Laboratory, Åbo Akademi University, a member of Turku Protein Core, FINStruct, and Biocenter Finland are gratefully acknowledged.

