## Supplementary figures for "LSD1 serine 166 is a phosphorylation switch for chromatin landscaping, gene activation, and tissue remodeling"

A

| Res. | Pos. | Sequence | PMID | Src | Cons. |
|---|---|---|---|---|---|
| T | 59 | EVGPGAVGERTPRKKEPPRAS | 18669648 | HTP | - |
| S | 80 | PPGGLAEPPGSAGPQAGPTVV | 18669648 | HTP | 0.59 |
| T | 104 | ATPMETGIAETPEGRRTSRRK | 18669648 | HTP | 0.48 |
| S | 126 | AKVEYREMDESLANLSEDEYY | 17081983 | HTP | 0.08 |
| S | 126 | AKVEYREMDESLANLSEDEYY | 18669648 | HTP | 0.08 |
| S | 131 | REMDESLANLSEDEYYSEEER | 15302935 | HTP | 0.21 |
| S | 131 | REMDESLANLSEDEYYSEEER | 17081983 | HTP | 0.21 |
| S | 131 | REMDESLANLSEDEYYSEEER | 18669648 | HTP | 0.21 |
| Y | 135 | ESLANLSEDEYYSEEERNAKA | 18669648 | HTP | 0.04 |
| Y | 136 | SLANLSEDEYYSEEERNAKAE | 18669648 | HTP | 0.04 |
| S | 137 | LANLSEDEYYSEEERNAKAEK | 15302935 | HTP | 0.13 |
| S | 137 | LANLSEDEYYSEEERNAKAEK | 17081983 | HTP | 0.13 |
| S | 137 | LANLSEDEYYSEEERNAKAEK | 18669648 | HTP | 0.13 |
| S | 166 | PQAPPEEENESEPEEPSGVEG | 15302935 | HTP | 1 |
| S | 166 | PQAPPEEENESEPEEPSGVEG | 17081983 | HTP | 1 |
| S | 166 | PQAPPEEENESEPEEPSGVEG | 18220336 | HTP | 1 |
| S | 166 | PQAPPEEENESEPEEPSGVEG | 18669648 | HTP | 1 |
| S | 849 | QATPGVPAQQSPSM | 18669648 | HTP | 0.78 |

C

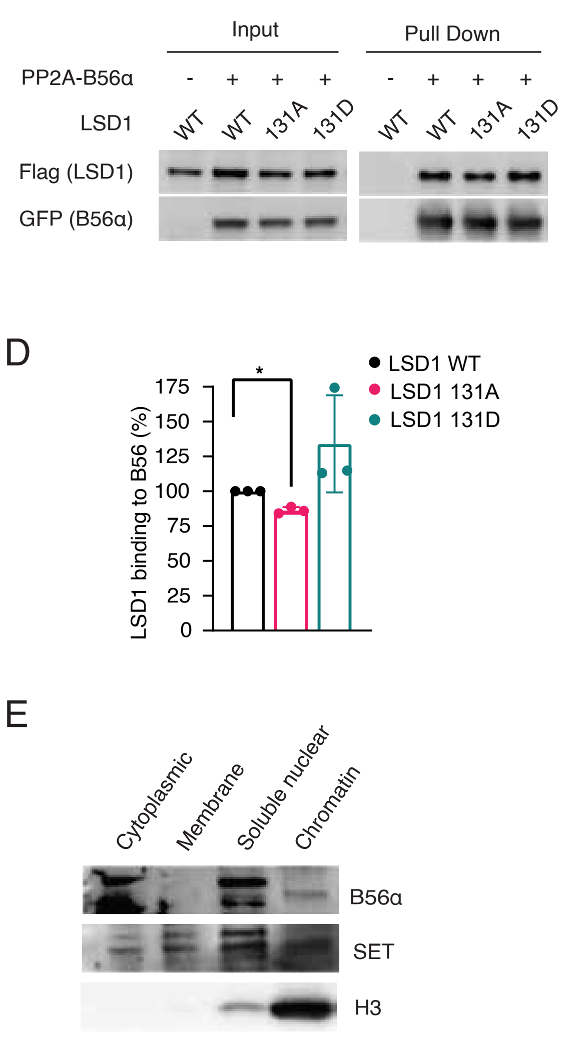


B

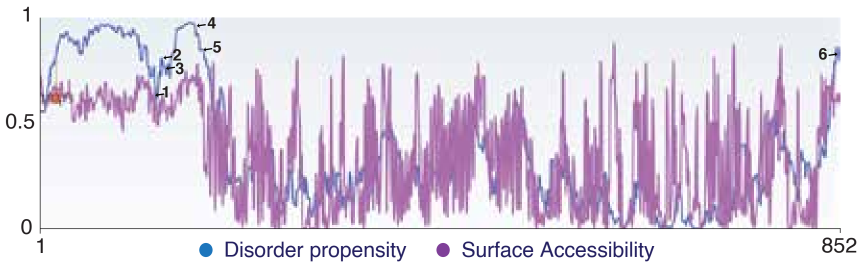


| No. | Res. | Pos. | Disorder | Accessibility |
|---|---|---|---|---|
| 1 | S | 126 | 0.6269 | 0.5836 |
| 2 | S | 131 | 0.8198 | 0.5868 |
| 3 | S | 137 | 0.7595 | 0.5229 |
| 4 | S | 166 | 0.9777 | 0.7321 |
| 5 | S | 172 | 0.8565 | 0.7276 |
| 6 | S | 849 | 0.8375 | 0.6218 |

F

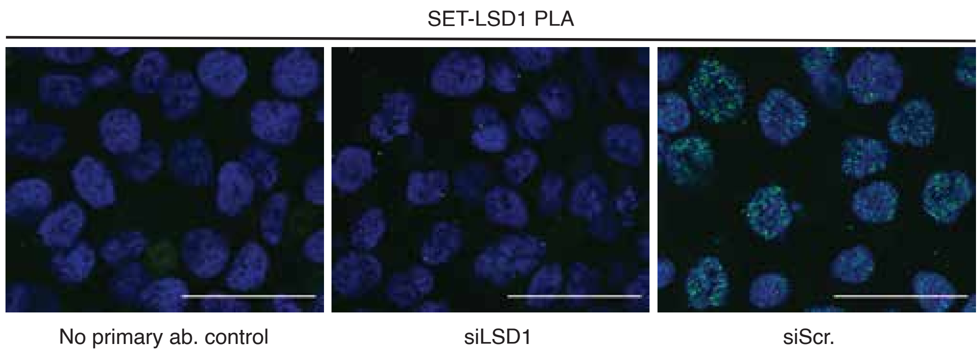


G

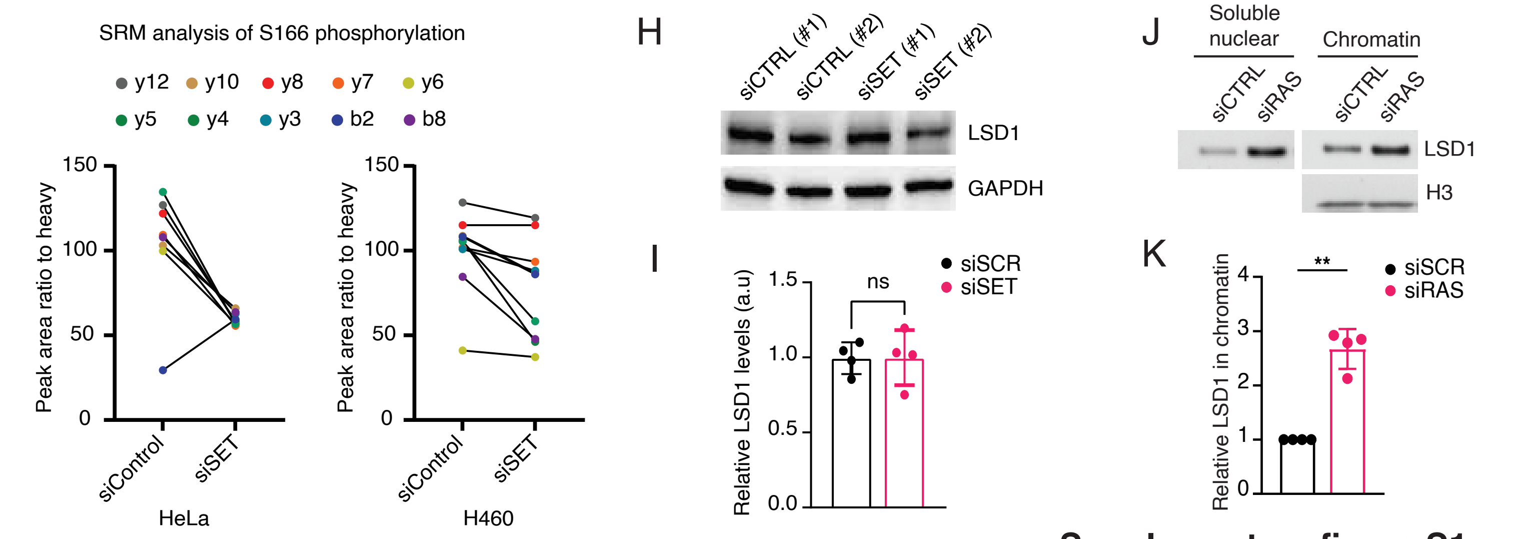


**Supplementary figure S1**

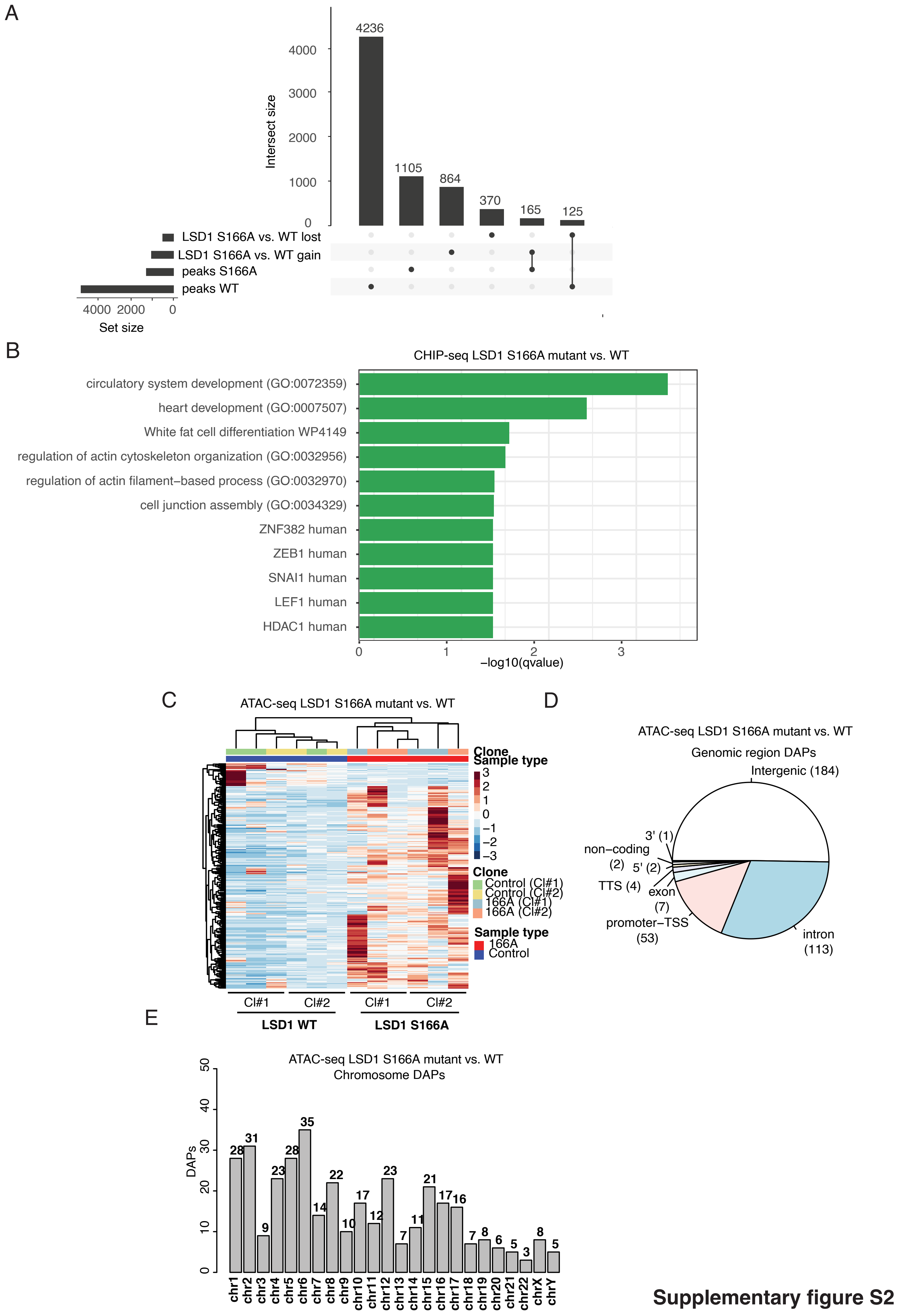


**Supplementary figure S2**

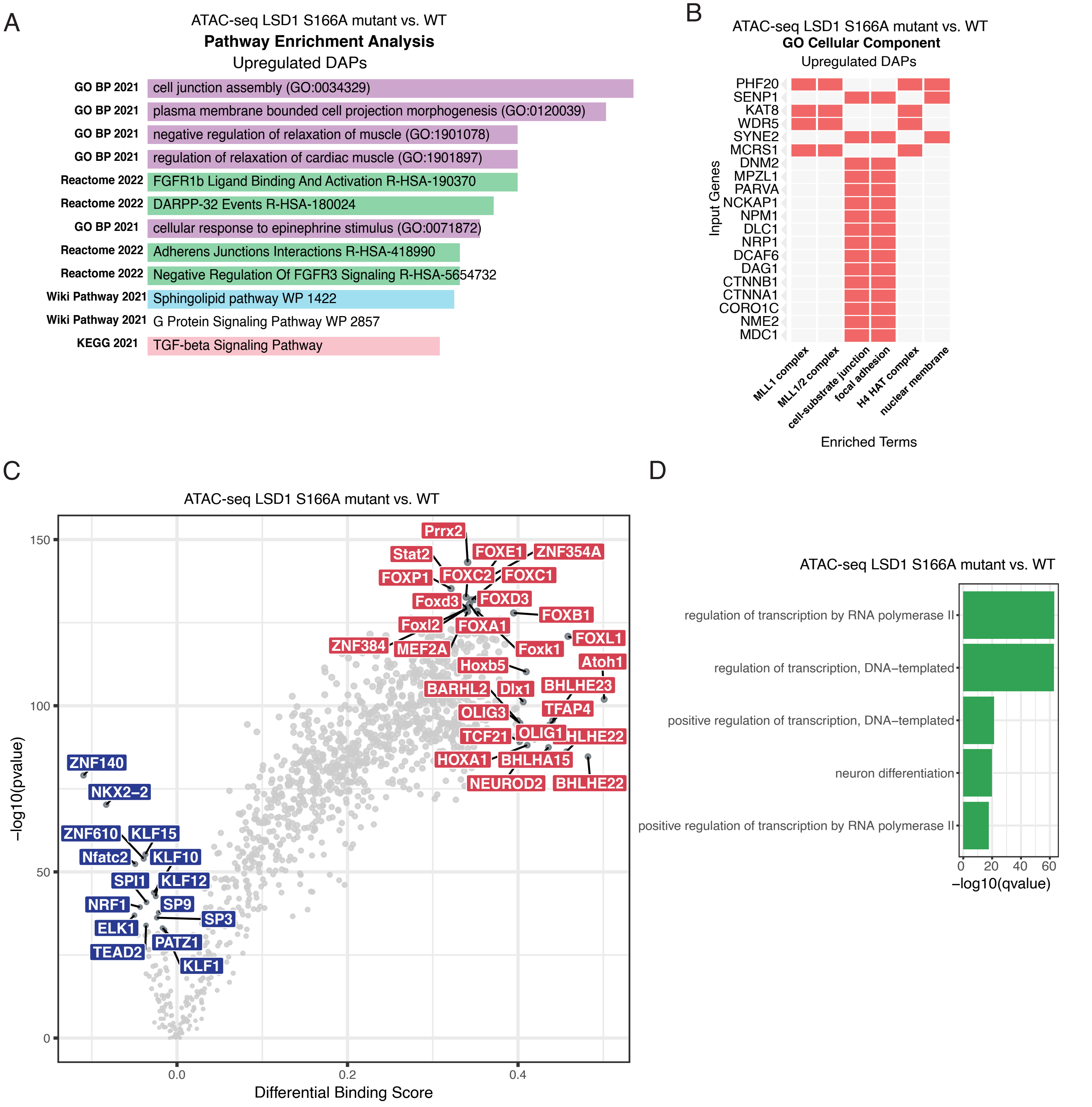


**Supplementary figure S3**

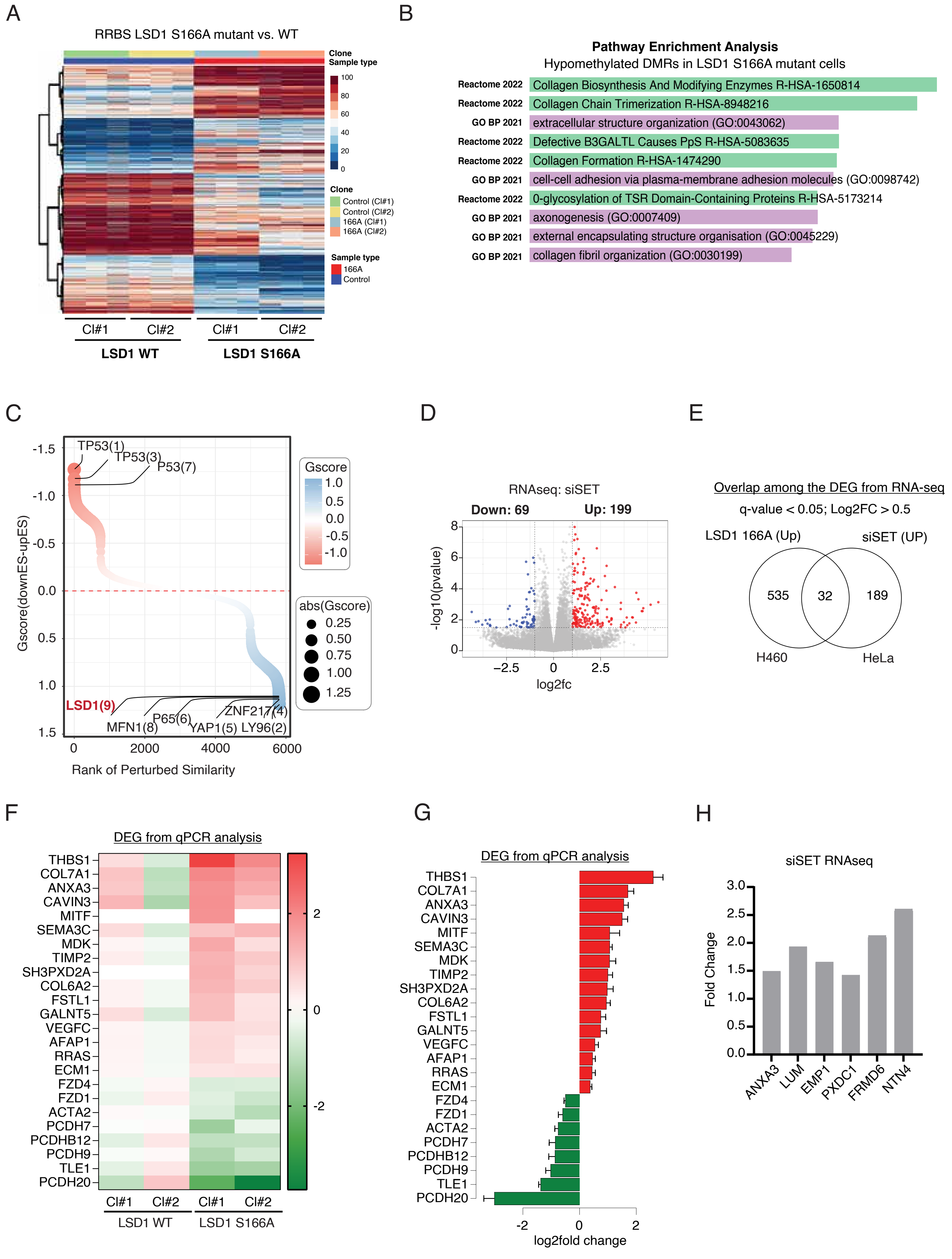


**Supplementary figure S4**

A

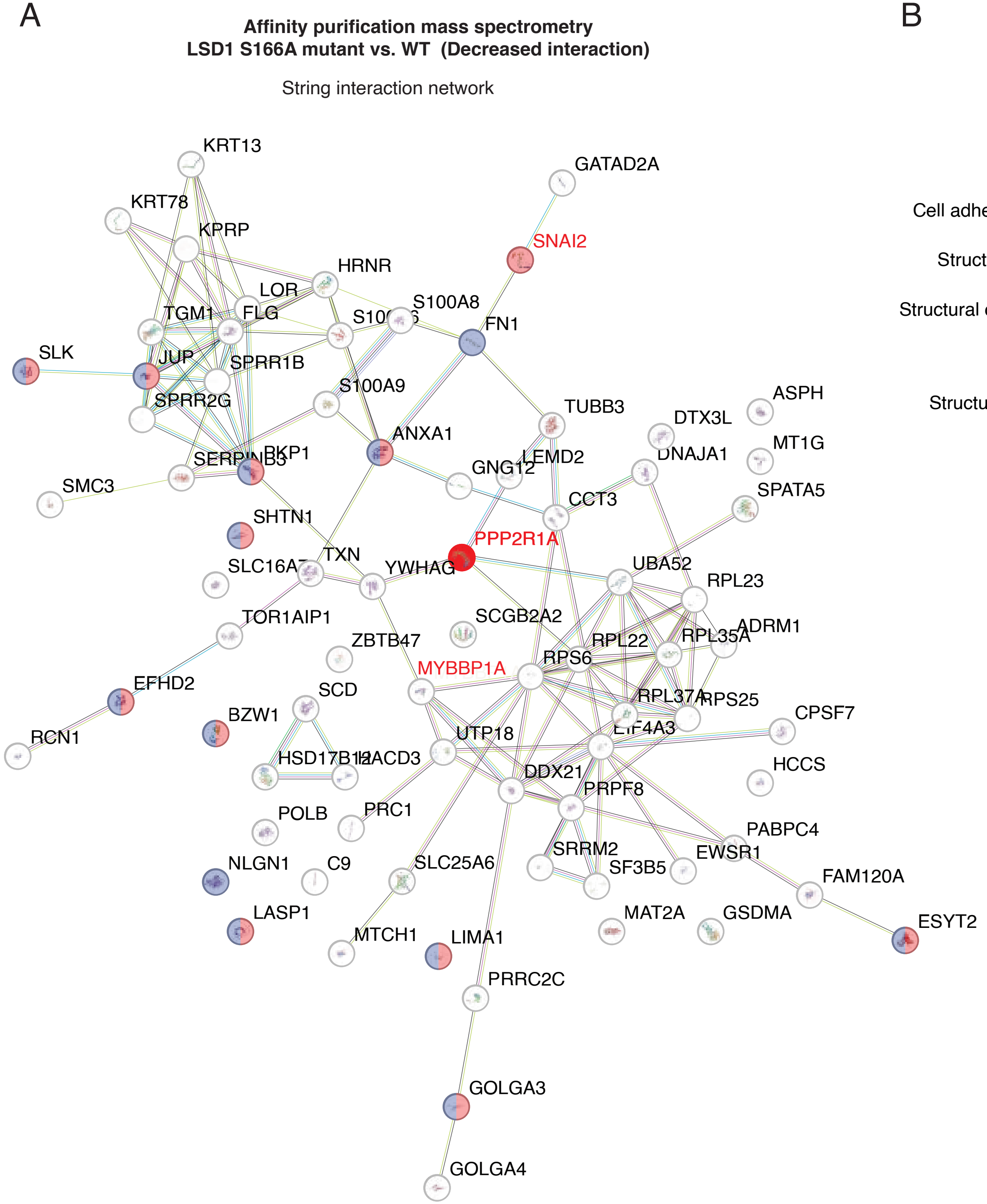


B

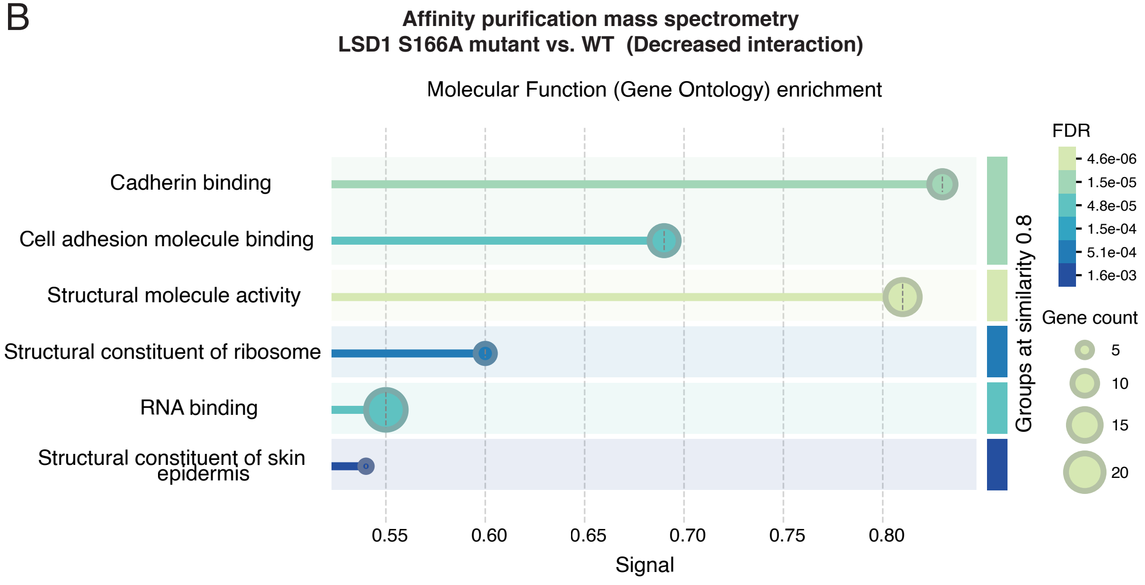


**Supplementary figure S5**

A

iPathway analysis based of DEGs

| Pathway name | Pathway Id | FDR |
|---|---|---|
| Viral protein interaction with cytokine and cytokine receptor[1] | 04061 | **3.449e-4** |
| ECM-receptor interaction[2] | 04512 | **4.168e-4** |
| Cytokine-cytokine receptor interaction[3] | 04060 | **4.404e-4** |
| Cell adhesion molecules[4] | 04514 | **4.404e-4** |
| Protein digestion and absorption[5] | 04974 | **4.404e-4** |

B

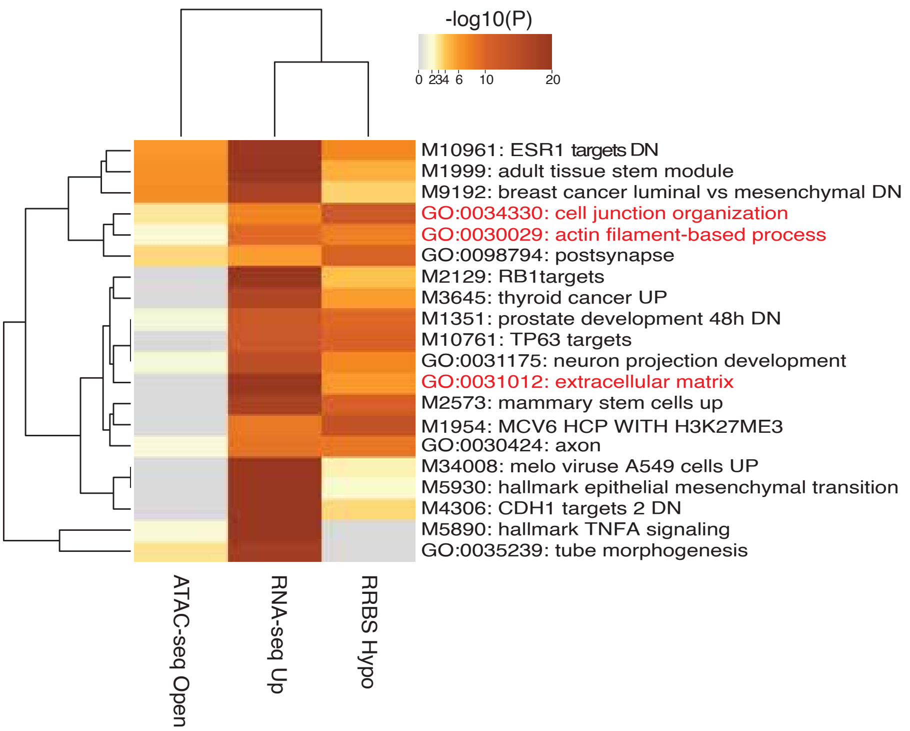


C

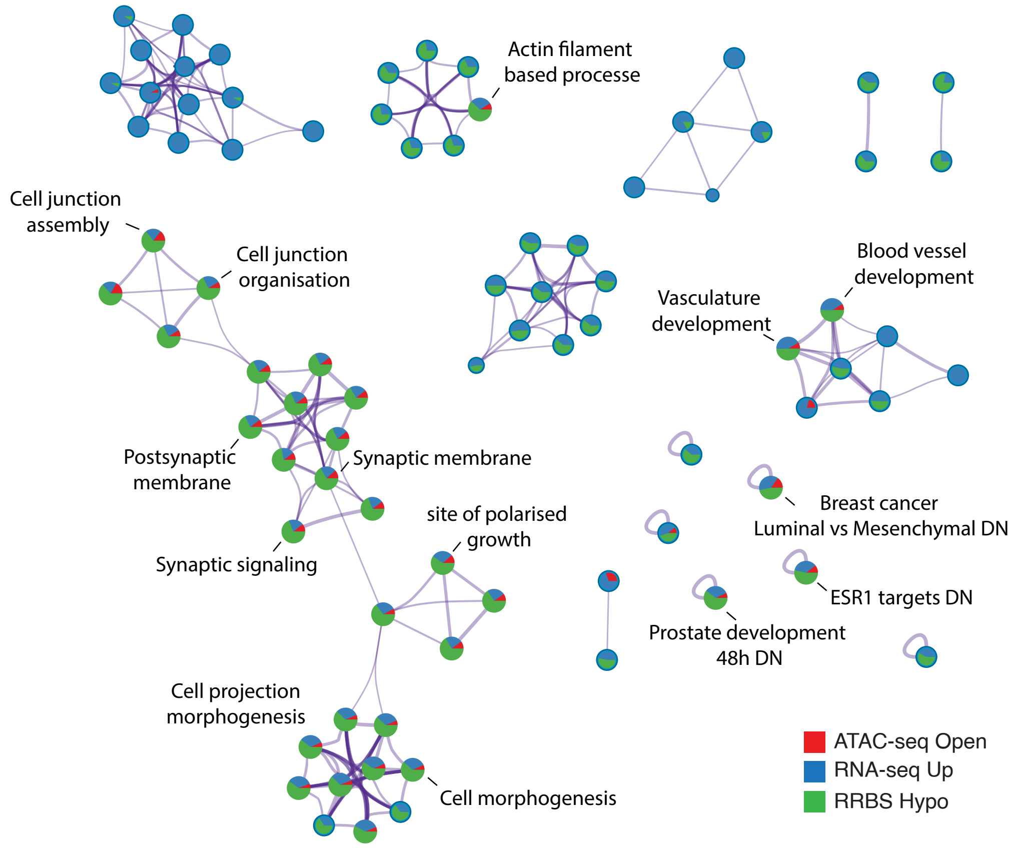


**Supplementary figure S6**

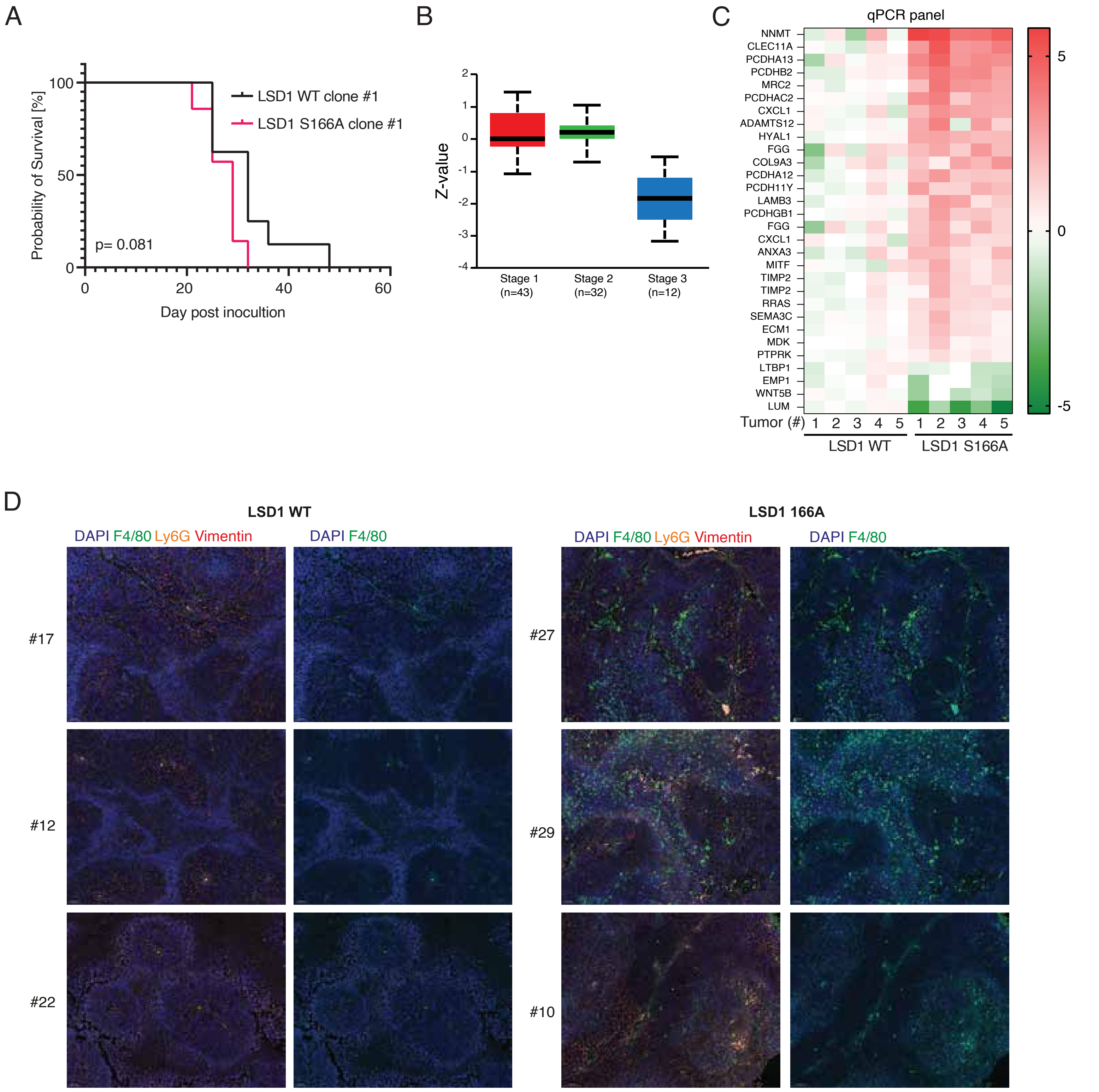


E

Top identified Oncogenic signatures gene sets.

| Gene Set | p-value | p-value (FDR) |
|---|---|---|
| BMI1 DN MEL18 DN.V1 UP | 5.673e-9 | 0.000001061 |
| BMI1 DN.V1 UP | 3.451e-8 | 0.000003226 |
| ERBB2 UP. V1 UP | 3.598e-7 | 0.00002243 |
| P53 DN.V1 DN | 5.435e-7 | 0.00002541 |
| MEL18 DN.V1 UP | 0.0000012 | 0.00004489 |

F

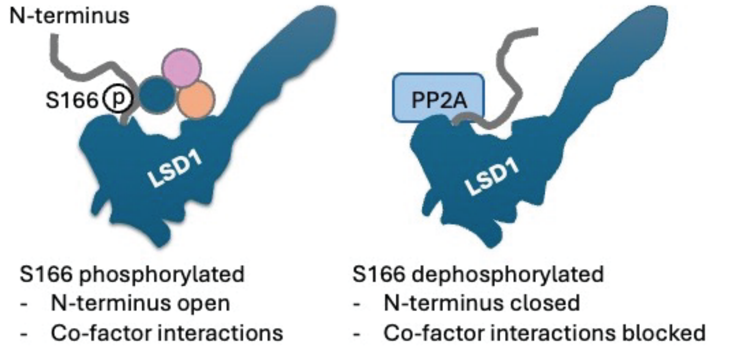


**Supplementary figure S7**
